# Spatial Biology and Organoid Technologies Reveal a Potential Therapy-Resistant Cancer Stem Cell Population in Pancreatic Ductal Adenocarcinoma

**DOI:** 10.1101/2025.05.22.655586

**Authors:** Pritha Adhikary, Jayati Chakrabarti, Saptarshi Mallick, Xi Sun, Ugonna Ezuma-Igwe, Abigail G. Branch, Jiang Wang, Nicolas Braissand, Megan P. Corbett, Sangmin Kim, Rohan Kanakamedala, Willian C. Bastian, Eugene Douglass, Meenu Priya Resmi, Payton Stevens, Marco Padilla-Rodriguez, Maga Sanchez, Gregory L. Beatty, Elliott B. Burdette, James Griffin, Petros G. Nikolinankos, Kellen Chen, Taylor Riall, Syed A. Ahmad, Rachna Shroff, Frédéric Hollande, Davendra Sohal, Yana Zavros

## Abstract

Pancreatic ductal adenocarcinoma (PDAC) remains one of the deadliest malignancies with a 5-year survival rate of less than 10%. Chemotherapy is the current standard-of-care (SOC) for advanced PDAC; however, treatment resistance driven by a complex and immunosuppressive tumor microenvironment (TME) limits its effectiveness. Cabozantinib (cabo), a multi-kinase inhibitor has shown promise in preclinical PDAC models and clinical trials, particularly when combined with immunotherapy, but its mechanisms of action within the PDAC TME remain unclear. Using CosMx™ Spatial Molecular Imaging and Orion™ Multiplex Immunofluorescence, we analyzed PDAC tissues collected from patients after first line systemic chemotherapy and surgery, or liver metastatic sites. Matched patient-derived organoid (PDO) co-cultures harboring cancer associated fibroblasts (CAFs), and autologous immune cells (IMM) (PDO/CAF/IMM) were generated and used to evaluate the effects of cabo and anti-PD1 immune check-point inhibitor pembrolizumab. Spatial analyses of post-chemotherapy and metastatic PDAC tissues revealed heterogeneous cell neighbor-hoods within the TME, including enrichment of Schwann cells, CAFs, T regulatory cells, and cancer stem cells (CSCs). Distinct niches were observed in metastatic liver tissues, characterized by CSCs expressing CD44 and TROP2. In PDO/CAF/IMM co-cultures, treatment with cabo in combination with pembrolizumab enhanced cancer cell death by depleting myeloid-derived suppressor cells and promoting cytotoxic T lymphocyte proliferation. Both patient tissues and treated co-cultures showed a persistent SOC-resistant CSC population that expressed CD44 variant 9 (CD44v9). This integrative spatial and organoid-based approach demonstrates that cabo potentially remodels the PDAC TME to potentiate PD-1 immunotherapy, while revealing therapy-resistant CD44v9+ CSCs as a potential therapeutic target.

## INTRODUCTION

Pancreatic ductal adenocarcinoma (PDAC) is among the most lethal cancers, with a 5-year survival rate of less than 10% despite advances in surgical and systemic therapies^1,2^. Most patients present with advanced or metastatic disease, and even those eligible for potentially curative resection face a high risk of relapse, with up to 80% experiencing local or distant recurrence within two years of diagnosis. Standard-of-care (SOC) chemotherapy regimens, such as FOLFIRINOX or gemcitabine/nab-paclitaxel, offer only limited survival benefits, underscoring the urgent need for new therapeutic strategies.

A major barrier to effective therapy in PDAC is the highly complex and immunosuppressive tumor microenvironment (TME), which consists not only of malignant epithelial cells but also cancer stem cells (CSCs), cancer-associated fibroblasts (CAFs), endothelial cells, immune cells, and neural elements^3,4^. Interactions among these diverse cell types contribute to profound tumor heterogeneity, facilitate immune evasion, and promote resistance to chemotherapy and immunotherapy. Although immune checkpoint inhibitors (ICIs) have revolutionized the treatment of several solid tumors, their efficacy in PDAC has been disappointing. This lack of response is thought to result from redundant and overlapping immunosuppressive mechanisms within the TME, including the infiltration of myeloid-derived suppressor cells (MDSCs) that inhibit cytotoxic T cell function, regardless of PD-1/PD-L1 blockade.

Cabozantinib is a tyrosine kinase inhibitor (TKI) targeting VEGFR (VEGF-1, VEGFR-2, and VEGFR-3), MET and AXL, and is approved as monotherapy for several solid tumors, including metastatic renal cell carcinoma^5,6^, hepatocellular carcinoma^7^, and medullary thyroid cancer^8^. By simultaneously inhibiting multiple signaling pathways including VEGFR, MET and AXL, cabozantinib disrupts tumor angiogenesis, proliferation, invasion, epithelial-to-mesenchymal transition, and metastasis^9–12^. While cabozantinib has demonstrated clinical benefit in other malignancies^5,6,7,8^, its efficacy in PDAC remains uncertain. Notably, a phase 1 clinical trial combining cabozantinib with gemcitabine failed to show meaningful benefit, in contrast to promising results from preclinical rodent models^13^. These discrepancies may reflect a limited understanding of therapy-resistant cell populations within the human TME and the potential inadequacy of traditional preclinical models to fully recapitulate the complexity of human PDAC.

Our previous work showed that cabozantinib can deplete polymorphonuclear MDSCs and, when combined with immune checkpoint blockade, induced tumor regression in preclinical PDAC models^14^. Our preclinical studies were the foundation for the development of an ongoing Phase II Trial evaluating the safety and efficacy of atezolizumab in combination with cabozantinib for the treatment of metastatic, refractory PDAC (NCT04820179). These findings, along with emerging evidence from clinical trials in other cancers^15–17^, provide a rationale for studying cabozantinib in combination with immune checkpoint inhibitors for PDAC.

In this study, we report the discovery of a persistent, SOC-resistant CSC population marked by the expression of the CD44 variant 9 (CD44v9) isoform. Our integrative approach provides new insights into the mechanisms of therapy resistance in PDAC and identifies CD44v9+ CSCs as a potential therapeutic target for overcoming relapse.

## MATERIALS AND METHODS

### Collection of patient biospecimens used for organoid and spatial biology analyses

Adult subjects 18 years of age or older having suspected and/or histologically or cytologically confirmed pancreatic cancer and willing and able to provide informed consent to undergo biopsy collection by an endoscopic procedure, and/or surgical tumor resection participated in these studies. Tumor tissues were collected by endoscopic ultrasound-guided fine-needle aspiration or core needle biopsies (University of Arizona Human Subjects Protection Program: IRB protocol number: 1912208231 (Zavros PI); IRB protocol number: 1099985869R001, TARGHETS) or during surgical resection (University of Arizona Cancer Center Biospecimen Repository IRB protocol number: 0600000609) to generate PDAC patient-derived organoids (PDOs). PDOs were also generated from core biopsies collected from patients with stage IV pancreatic adenocarcinoma, confirmed by histology or cytology, and willing and able to provide informed consent to be enrolled in the Phase II Atezolizumab + Cabozantinib in Patients w/ Metastatic, Refractory Pancreatic Cancer clinical trial (ClinicalTrials.gov ID NCT04820179, IRB protocol number: 2108097176). All biospecimens were de-identified prior to transfer to the Zavros laboratory for analyses and generation of PDOs. Commercially available tissue microarray slides of patient normal pancreas and PDAC biospecimens was used for the spatial transcriptomic analyses and included TissueArray.com, catalogue numbers

TissueArray.com (PA242e): https://www.tissuearray.com/tissue-ar-rays/Pancreas/PA242e?srsltid=AfmBOord_2o_bfyH09yMTMc86Dsg1svJtloNcxP4jTVUg_O7NqLEkU77

TissueArray.com (PA242f): https://www.tissuearray.com/tissue-ar-rays/Pancreas/PA242f?srsltid=AfmBOopYHVF0gBrWkbpGh_hJR58AbNXS50nDRRmGJEwe7TV1V6Fw747C. **Supplemental Table 1** details the patient biospecimen characteristics from which the organoid lines and spatial biology analyses were performed.

### Generation and culture of human PDAC Patient Derived Organoids (PDOs) from tumor tissues

Patient derived organoids from tumor tissues and biopsies collected from patients with PDAC were generated using a protocol published by our research team ^14,18^. Human tumor tissues and biopsies were washed in Dulbecco’s phosphate-buffered saline without calcium and magnesium (DPBS, Thermo Fisher Scientific, 14190-144) supplemented with 1% Penicillin/Streptomycin (Thermo Fisher Scientific, 15-140-122), 1% Kanamycin Sulfate (Thermo Fisher Scientific, 15-160-054) and 0.2% Gentamicin/Amphotericin B (Thermo Fisher Scientific, R-01510). Tissue was then dissected into approximately 1-2 mm pieces and enzymatically digested using 5mL Hank’s balanced salt solution (HBSS, Thermo Fisher Scientific, 14175095) supplemented with Collagenase P (1mg/mL, SIGMA, 11 213 865 001), 5% FCS (FCS, R&D System, S12450H), 1% Penicillin/Streptomycin, and 1% Kanamycin Sulfate for 15 minutes at 37°C with gentle agitation/shaking. After digestion, an equal volume of HBSS supplemented with 5% FCS, and 1% Penicillin/Streptomycin, 1% Kanamycin Sulfate and 0.2% Gentamicin/Amphotericin B was added, and dissociated tissue was centrifuged for 5 minutes at 400xg. The cell pellet was washed with DPBS, resuspended in the appropriate volume of Matrigel^TM^ (Corning, 356231), seeded in domes in tissue culture plates (MIDSCI, 92024) and/or glass bottomed chambered slides (Cellvis, C815.HN). Matrigel^TM^ domes reconstituted with tumor cells formed a solid gel after 20-25 minutes at 37°C and were then overlaid with organoid media containing DMEM/F12 (Thermo Fisher Scientific, 12634010), 1x B27 (Thermo Fisher Scientific, 12587010, 284μM ascorbic acid (R&D Systems, 4055-50), 20μg/mL Insulin (R&D Systems, 3435/10), 0.25μg/mL hydrocortisone (SIGMA, H0888), 100ng/mL FGF2 (Peprotech, 100-18B), 100nM ATRA (SIGMA, R2625), 10μM Y27632 (SIGMA, Y0503), 100ng/mL FGF10, 1% Penicillin/Streptomycin, 0.1% Gentamicin/ Amphotericin B, 2mM Glutamax (Fisher Scientific, 350-50-061), and 56μg/mL Bovine Pituitary Extract (Thermo Fisher Scientific, 13028014). Cultures were maintained at 37°C in 5% CO_2_ and media replenished every 3-4 days.

Approximately 7-10 days post-seeding, PDOs were harvested in ice cold DPBS without calcium and magnesium and centrifuged at 400xg for 5 minutes 1-2 times to remove Matrigel^TM^. Supernatant was removed and PDO pellet was resuspended in ACCUTASE™ Cell Detachment Solution (STEMCELL Technologies, 07922) and stored for 5-10 minutes at 37°C in a 5% CO_2_ incubator. After incubation, PDOs were gently passed through a 26-gauge needle (BD, 309597), washed with DPBS supplemented with 1% Penicillin/Streptomycin and 1% Kanamycin Sulfate, resuspended in Matrigel^TM^ domes and seeded in culture dishes and/or glass bottom chambered slides as described for PDO/CAF/immune cell co-cultures and analyses. Expanded PDO cultures were also harvested and cryopreserved using CryoStor® CS10 (STEMCELL Technologies, 07930) according to the manufacturer’s protocol.

### Isolation and culture of Peripheral Blood Mononuclear Cells (PBMCs) and immune cells from patient whole blood

#### Peripheral Blood Mononuclear Cells (PBMCs)

Peripheral Blood Mononuclear Cells (PBMCs) were cultured from autologous whole blood collected from patients with pancreatic ductal adenocarcinoma from which PDOs were generated. Lymphoprep^TM^ density gradient medium was used for the isolation of mononuclear cells from whole blood according to the manufacturer’s protocol (STEMCELL Technologies, catalogue number 07851). Briefly, blood was diluted with DBPS containing 1% FCS and 1% Penicillin/Streptomycin, slowly loaded onto the SepMate^TM^ 50mL tube containing 15mL of Lymphoprep^TM^ and centrifuged for 10 minutes at 1200 x g. Leukocytes were collected, diluted with DBPS containing 1% FCS and 1% Penicillin/Streptomycin and centrifuged at 300 x g for 8 minutes at 4°C. Isolated PBMCs were then resuspended with DBPS containing 1% FCS and 1% Penicillin/Streptomycin and centrifuged at 120 x g for 10 minutes at 4°C to wash. The PBMCs were then used for cultures of immune cells as follows and published by our research team ^14^:

#### Dendritic Cells (DCs)

Approximately 1 million PBMCs were seeded into each well of a 24 well plate. AIM V dendritic cell culture medium containing AIMV medium (Thermofisher Scientific, 12055-091) supplemented with 10% human serum AB (Gemini Bio, 800-120), 50 μM β-mercaptoethanol (Thermo Fisher Scientific 21985023), 800 U/mL granulocyte-macrophage colony-stimulating factor (GM-CSF, Thermo Fisher Scientific, PHC6025) and 500 U/mL interleukin 4 (IL-4, Thermo Fisher Scientific, RIL4I) for 3 consecutive days. Cultures were then maintained for 24 hours using AIM V dendritic cell culture medium detailed above with the addition of 5 ng/mL tumor necrosis factor α (TNF-α, Thermo Fisher Scientific, BMS301), 5 ng/mL interleukin 1β (IL-1β, Thermo Fisher Scientific, RIL1BI), 150 ng/mL interleukin 6 (IL-6, Thermo Fisher Scientific, RIL6I) and 1 μg/mL prostaglandin E2 (PGE2, TOCRIS, 2296) for maturation.

#### Cytotoxic T Lymphocytes (CTLs)

Cytotoxic T Lymphocytes (CTLs) were isolated and cultured from PBMCs using the EasySep^TM^ Human CD8+ T cell Enrichment Kit (STEMCELL Technologies, 19053) following the manufacturer’s protocol. Briefly, 1 million PBMCs were incubated for 10 minutes at room temperature with 50 μL/mL Enrichment Cocktail followed by a 5-minute incubation with the 150 μL/mL Magnetic Particles diluted in EasySep Buffer (STEMCELL Technologies, 20144). CD8+ T cells were then magnetically separated using the EasySep Magnet (STEMCELL Technologies, 18000). Supernatant containing CD8+ T cells was then transferred to a separate tube and centrifuged at 300 x g for 5 minutes. Isolated CD8+ T cells were cultured using CTL culture media (RPMI 1640 medium (Thermo Fisher Scientific, 11875-093) containing 10% human serum AB (Gemini Bio, 800-120), 1% penicillin/streptomycin (Fisher Scientific, 15-140-122), 50μM β-mercaptoethanol (Thermo Fisher Scientific, 21985023), 1x insulin-transferrin-selenium (Thermo Fisher Scientific, 41400045), 0.15μg/mL interleukin 2 (IL-2, Thermo Fisher Scientific, RP-8608) and 0.1μg/mL interleukin (IL-7, Thermo Fisher Scientific, RIL75)).

#### Myeloid-Derived Suppressor Cells (MDSCs)

Myeloid-Derived Suppressor Cells (MDSCs) were isolated and cultured from PBMCs using monocyte culture medium containing AIM V medium supplemented with 10 ng/mL Interleukin-1 beta (IL-1β, Thermo Fisher Scientific, 12055091), 10 ng/mL Interleukin-6 (IL-6, Thermo Fisher Scientific, RIL6I), 1 μg/mL Prostaglandin E2 (PGE2, TOCRIS, 2296), 2 ng/mL Transforming Growth Factor Beta 1 (TGF-β1, R&D Systems, 7754-BH/CF), 10 ng/mL Tumor Necrosis Factor alpha (TNF-α, Thermo Fisher Scientific, BMS301), 10 ng/mL Vascular Endothelial Growth Factor (VEGF, Thermo Fisher Scientific, RVEGFI) and 10 ng/mL Human Granulocyte-Macrophage Colony Stimulating Factor (GM-CSF, Thermo Fisher Scientific, PHC6025) and 50% conditioned medium collected from actively growing autologous PDO cultures.

### Establishment, experimental design and image analyses of autologous Patient Derived Organoid (PDO)/cancer associated fibroblast (CAF)/Immune Cell (IMM) co-cultures

Co-cultures were established using isolated DCs, CTLs and MDSCs following our published protocols ^14^. After maturation, DCs were pulsed for 2 hours using conditioned medium collected from autologous and actively growing PDOs. Matured and pulsed DCs were then co-cultured with CD8+ CTLs for 24 hours. PDOs were harvested following the described passaging protocol above without ACCUTASE™ Cell Detachment Solution dissociation. Co-cultures were established using approximately 70-80% PDO confluency and IMM cell ratios of 4:1:4 (DC:CTL:MDSC) and maintained in PDO culture medium at 37°C in 5% CO_2_ for 24-48 hours prior to treatment under the following experimental conditions: 1) PDOs vehicle, 2) PDOs+CTLs vehicle, 3) PDOs+CTLs 10 μM Cabozantinib, 4) PDOs+CTLs 0.5 μg/mL Pembrolizumab, 5) PDOs+CTLs+MDSCs vehicle, 6) PDOs+CTLs+MDSCs 10 μM Cabozantinib, 7) PDOs+CTLs+MDSCs 0.5 μg/mL Pembrolizumab, and 8) PDOs+CTLs+MDSCs 10 μM Cabozantinib plus 0.5 μg/mL Pembrolizumab.

PDO/CAF/IMM co-cultures were imaged every 24 hours over 3 consecutive days under Phase 10X using the Nikon Spinning disk Ti2 Eclipse inverted confocal microscope (Nikon Corporation, Tokyo, Japan). As a measure of cell death, treated co-cultures actively growing between days 3-5 were treated with propidium iodide (Biolegend 421301) 30 minutes prior to image analysis. Nikon NIS Elements AR 5.42.03 software with the General Analysis 3 (GA3) module was used for image processing and analysis of the co-culture samples. For the DAPI and PI channels, the following image processing tools were used to achieve accurate segmentation: Rolling ball background subtraction (Radius = 10μm) and Local Contrast (Degree = 50%; Radius = 8.00μm). For the Phase contrast channel, the following image processing tools were applied: Low Pass Filter (Level/strength = 30px), Detect Edges, Local Contrast (Degree = 75%; Radius = 120.00μm), and Gamma Correction (gamma = 0.8). Using the Bright Spots Detection tool, the DAPI and PI channels were maintained at threshold using the following parameters: Diameter = 6 μm, Contrast = 0.010, Symmetry = all objects, and the same intensity threshold for the PI channel for across all images. A Signal Intensity threshold was used to detect and segment the organoids in the Phase-contrast channel. The Dilate and Close Hole tools were used to mask the entire organoid to achieve accurate quantification of organoid circumference. An Area Filter was applied to remove small, false positive organoid detections. The AND binary operation was performed between DAPI/PI channels and corresponding organoid (Phase-contrast) channel to restrict analysis only to cells present in detected organoids. For each organoid, the following parameters were quantified: Circumference, Circularity, and DAPI and PI object count (normalized to 100μm of organoid circumference).

### Isolation and culture of PDO-generated cancer associated fibroblasts (CAFs)

Culture plates (24 wells) were coated using a 1:40 dilution factor of Matrigel^TM^ to Advanced DMEM-F12 medium for 16 hours at 37°C in a 5% CO_2_ incubator. Matrigel^TM^ was then removed and immediately used for generation of monolayers and CAF cultures. PDOs were harvested in cold DPBS and centrifuged at 400xg for 5 min at 4°C to remove the Matrigel^TM^ and resuspended in 2 mL 0.25% Trypsin-EDTA supplemented with 10 µM Y-27632 (ROCK inhibitor) for 5–7 minutes at 37°C in 5% CO₂. Following enzymatic digestion, organoids were mechanically dissociated by pipetting 10–12 times to obtain a single-cell suspension. The enzymatic reaction was neutralized by adding Advanced DMEM/F-12 supplemented with 10% FCS and cells were subsequently centrifuged at 400xg for 5 minutes. The isolated cell pellet was resuspended in DMEM/FCS medium (Advanced DMEM/F-12 supplemented with 10% FCS and 1% Penicillin-Streptomycin), seeded onto the prepared Matrigel^TM^-coated plates and cultured at 37°C in 5% CO₂. The culture media was removed and replenished every alternate day. Once the monolayers reached approximately 80-90% confluency, cells were passaged using 0.25% Trypsin-EDTA supplemented with 10 µM Y-27632, as described above, and replated onto freshly coated plates for four subsequent passages to selectively enrich for CAFs/fibroblasts.

### Spectral flow cytometry (Cytek™ Aurora)

The Cytek*™* Aurora advanced spectral flow cytometry system was used to analyze changes in cancer associated fibroblast phenotype, tumor cell and immune cell viability and proliferation in response to Cabozantinib and immune checkpoint inhibition within the human PDO co-cultures and mouse orthotopic transplants. This approach allowed us to use a unique combination of 16 fluorochromes simultaneously in one sample to achieve single cell resolution multiplexed data using the PDO/CAF/IMM cell co-cultures **(Supplemental Table 2)**. The multicolor flow cytometry panel was designed using the Cytek®Full Spectrum Viewer to calculate the similarity index (**Supplemental Figure 1**). PDO co-cultures were harvested in cold serum free organoid media, centrifuged at 400 x g for 5 minutes and dissociated to single cells using ACCUTASE™ (Thermo Fisher Scientific, 00-4555-56). Cells were then washed by centrifugation at 400xg for 5 minutes and incubated with fluorochrome-conjugated/unconjugated primary surface antibodies at 4°C for 30 minutes. Cells were washed with Cell Staining Buffer (BioLegend catalogue no: 420-201) and incubated with secondary antibody when required at 4°C for 30 minutes (**Supplemental Table 2**). Cytoplasmic staining was performed by first incubating cells with Cytofix/Cytoperm™ Fixation/Permeabilization Solution (BD Biosciences, 554714) at 4°C for 20 minutes, followed by washing with Fixation/Permeabilization wash buffer. Cells were then labeled with fluorochrome-conjugated/unconjugated intracellular/cytoplasmic primary antibodies at 4°C for 30 minutes, washed and incubated with secondary antibodies when needed at 4°C for 30 minutes (**Supplemental Table 2**). Cells were resuspended in cell staining buffer in a volume of 150 μl and fluorescence intensity measured using the Cytek Aurora 5 Laser Spectral Flow Cytometer. Unstained cells were fixed and used as reference controls. UltraComp eBeads™ Compensation Beads (Thermo Fisher Scientific, 01-2222-42) were stained with the individual antibodies and used as single stain control for compensation and gating. Data was acquired using the Cytek™ Aurora and analyzed using Cytobank software (Beckman Coulter, Indianapolis, IN) or FlowJo software (BD).

### Nuclear Morphometric Analysis (NMA)

Cell viability was calculated as a measure of nuclear morphology using the Nuclear Morphometric Analysis (NMA) published protocol ^19^. PDO nuclei were captured using the Zeiss LSM 880 confocal microscope Nikon Spinning disk Ti2 Eclipse inverted confocal microscope (Nikon Corporation, Tokyo, Japan). Images were imported into the ImageJ Nuclear Irregularity Index (NII) plugin for the measurement of key parameters that included cell area, radius ratio, area box, aspect, and roundness. Using the published spreadsheet template containing formulas for measurements ^19^, the NII of each cell was calculated as: *NII = Aspect – Area Box + Radius Ratio + Roundness*

The distribution of area vs NII of vehicle-treated cells were represented on a scatter plot and set as normal cell nuclei. The same scatter plots were generated for each experimental condition, and the NII and area of treated cells were compared to the normal nuclei, and classified as one of the following NMA populations: Normal (N: normal shape and size; without damage that affects nuclear morphology), Irregular (I: normal size with high irregularity; mitotic catastrophe), Small Regular (SR: condensed and regular morphology; apoptosis), Small (S; spherical and small, mitosis), Small Irregular (SI: condensed and small irregular morphology; mitosis with damage), Large Regular (LR: large and regular morphology; senescence), or Large Irregular (LI: significant nuclear damage in large nuclei with muti-nucleated cell morphology; mitotic catastrophe). Cells classified as SR exhibited early stages of apoptosis, and cells classified as either I, SI or LI exhibited significant nuclear damage. The percentage of cells in each NII classification category were calculated and plotted as a histogram using GraphPad Prism ^19^.

### Human tumor tissue histology and immunofluorescence staining

Human tumor tissues were formalin fixed (3.7%), paraffin embedded (FFPE) and serially sectioned at a 5-micron thickness. After deparaffinization and antigen retrieval (Antigen Unmasking Solution, Vector Laboratories H-3300-250), slides were blocked with 20% normal donkey serum for 20 min at room temperature and incubated with primary antibodies overnight at 4°C. Combinations of the following primary antibodies were used: CD44v9 (Cosmo Bio, LKGM001, 1:1000), Amylase (Invitrogen, PA5-25330, 1:100), and cytokeratin 19 (CK19, NB1-85601, 1:100). After primary antibody incubation, slides were washed with 0.01% Triton X-100/PBS and then incubated with AlexaFluor^TM^ secondary antibodies (Thermo Fisher Scientific) including anti-rat AlexaFluor^TM^ 488 (A-21208, 1:100), anti-rabbit Alexa Fluor^TM^ 647 (A-31573), anti-mouse Alexa Fluor 555 (A-31570) at room temperature for 1 hour. Nuclei were counterstained with Hoechst 33342 (Thermo Fisher Scientific, H1399, 1:1000). The slides were mounted using Fluoromount-G^TM^ (Invitrogen, 00-4958-02) and images captured using Nikon Spinning disk Ti2 Eclipse inverted confocal microscope.

### Whole mount immunofluorescence of PDAC PDO cultures

Whole-mount immunofluorescence staining of PDAC PDO cultures was performed as published by our laboratory ^20^. Cultures grown and experimentally treated in 8 well glass bottom chamber slides (Cellvis, C8-1.5H-N) were fixed using 3.7% formaldehyde (ThermoFisher Scientific, 28908) at room temperature for 15 minutes or at 4°C for 12-16 hours and then permeabilized with 0.5% Triton X-100/PBS (Millipore Sigma, X100) at room temperature for 20 minutes. Proliferation was measured using the Click-iT™ EdU Cell Proliferation Kit for Imaging, Alexa Fluor™ 555 dye (ThermoFisher Scientific, C10338) according to the manufacturer’s protocol. In a separate series of experiments, slides were blocked using 2% donkey serum (Jackson ImmunoResearch, 017-000-121) at room temperature for 1 hour followed by incubation with primary antibodies at 4°C for 12-16 hours. Primary antibodies used for the studies included: human anti-CD44v9 (Cosmo Bio, LKGM001, 1:1000) and anti-fibronectin (Invitrogen, 14-9869-82, 1:1000). After primary antibody incubation, organoids on slides were washed with 0.01% Triton X-100/PBS and incubated with Alexa Fluor secondary antibodies (ThermoFisher Scientific) including anti-rat AlexaFluor^TM^ 488 (A-21208, 1:100) and anti-mouse Alexa Fluor 555 (A-31570) at room temperature for 1 hour. Nuclei were counterstained with Hoechst 33342 (ThermoFisher Scientific, H1399, 1:1000). Images were captured using the Nikon Ti2-E Inverted Microscope (with a Crest X-Light V2 L-FOV Spinning Disk Confocal). Automated measurement of fluorescence intensity was calculated using the Nikon Element Software Version 5.21.02.

### CosMx^TM^ Spatial Molecular Imaging (SMI) sample preparation and workflow

The CosMx™ SMI’s unique combination of high-plex, high-throughput single cell spatial profiling was used to identify inter- and intra-patient differences within the PDAC tumor microenvironment. Patient formalin fixed paraffin embedded (FFPE) tissue sections were analyzed by NanoString CosMx™ SMI using tissue preparation and workflow according to the manufacturer’s protocol (Nanostring, MAN-10186-01-1). The CosMx SMI instrument was used for slide scanning, FOV selection, and automated hybridization chemistry to detect target molecules/genes (1000 plex). Briefly, PDAC tissues were formalin fixed (3.7% formalin) and paraffin embedded (FFPE) and sectioned at a 5μm thickness. Sections were then permeabilized and RNA probes were hybridized to their targets in the tissue sample. The tissue sample is washed, then incubated with oligolabeled antibodies for morphology marker staining (DAPI Nuclear Stain, UV channel, CD298/B2M Segmentation Marker Mix Yellow channel, PanCK Marker Mix Blue channel, CD45 Marker Mix Red channel). After washing, the flow cell was assembled and loaded onto the SMI instrument for morphology marker imaging. The desired imaging area(s) or fields of view (FOVs) on the tissue (up to 100 mm^2^) were selected. The instrument then automated rounds of reporter binding and fluorescent imaging to read out the barcodes on each imaged RNA probe or protein antibody. Data acquisition was completed on the instrument and all data storage and analysis was completed within the cloud-based account. SMI provided a high-resolution subcellular image-based readout by identifying the X, Y, and Z coordinates of each target gene, which is translated to spatial location data and exported to the cloud. The optical system includes an epi-fluorescent configuration that uses custom water immersion optics with 1.1 NA and 22.77X magnification. The FOV size is 0.7mm x 0.9 mm. Illumination was widefield with a mix of lasers and LEDs to allow for UV cleavage (385 nm) and imaging of four different fluorophores that included Alexa Fluor-488, Atto-532, Dyomics Dy-605, and Alexa Fluor-647.

### CosMx^TM^ Spatial Molecular Imaging (SMI) analysis

Raw expression matrices and FOV segmentation data of FFPE sections of tissues obtained from the CosMx SMI were analyzed in R 4.5.1 using the Seurat v5.5.1 package^21^. Expression matrices were read using Seurat’s LoadNanostring function to create a basic Seurat object, used for further processing and analysis. Metadata obtained from the instrument was applied to the object during this step. Quality control and filtering were performed based on unique feature count, number of transcripts, cell area, and negative probe fraction metrics. Seurat objects of samples from different slides were merged into a single object using Harmony integration^22^, to account for batch effects and allow substantive downstream comparative analyses. The combined object was then processed using Seurat’s standard clustering pipeline: SCTransform normalization and regularization was applied to the 1000 plex dataset and log-normalization to the whole-transcriptome dataset, identification of principal components, UMAP generation, and finally Louvain clustering for an unsupervised identification of cell types. Following processing, cell clusters were assigned cell types based on known gene signatures and published data ^23–27^, and supervised identification using the scType package ^28^. Cell types and associated lists of marker genes were compiled based on various published studies. Spatial niches were identified using Seurat’s BuildNicheAssay function with niches.k set to 7. Analyses performed using the processed Seurat object included identification of key differentially expressed genes using FindAllMarkers, violin plots for specific gene features using VlnPlot() together with Wilcoxon tests to derive p-values, heatmaps comparing overall gene expression profiles among patients and cell type clusters using DoHeatmap, dot plots contrasting gene expression between groups for specific sets of marker genes using DotPlot, and feature plots displaying the expression of specific genes across identified cell clusters using FeaturePlot. The data was simultaneously analyzed using Seurat’s set of spatial analysis tools: spatial UMAP (ImageDimPlot), spatial feature plot (ImageFeaturePlot), spatial niches (ImageDimPlot colored by niche) and isolation of FOVs of interest for comparison and differential gene expression profiling. Within each sample, FOVs were compared using the spatial image plot and differential expression of genes of interest between tumor and adjacent normal sections of the tissue. Stacked bar plots showing distribution of clusters across niches and tissues were generated with the ggbar() function from the ggplot2 package.

Signaling interactions between spatial niches and cell clusters within each niche were identified using CellChat v2.1.0, using the standard analysis pipeline (subsetData, identifyOverExpressedGenes, identifyOverExpressedInteractions, computeCommunProb) and ligand-receptor database v2 with preprocessing to convert CosMx SMI spatial coordinates to a format recognized by CellChat. Enriched pathways relevant to the identified interactions were determined using CellChat’s computeCommunProbPathway function and visualized in a heatmap format using netAnalysis_signalingRole_heatmap on the ‘netP’ slot of the CellChat object. Network graphs were generated by exporting the net interaction weights from CellChat objects and filtering the matrix to include interactions involving a specific cell cluster. The matrix was then used to construct an adjacency list representation, modeling a graph where each edge represented a directed interaction from one cell cluster to another with weight equal to interaction strength and vertices representing cell clusters. The adjacency list was visualized using the qgraph v1.9.8 package in R.

Nearest-neighbor proximity analysis was computed by filtering gene expression positive cells as the query, and different groups of cells as the target. For each comparison, a nearest-neighbor analysis was conducted by running get.knnx from the FNN v1.1.41 package in R. The distance of each pair was then quantified by converting pixels to microns with a conversion factor of 0.18 µm/px, and a histogram was generated using gghistogram from ggpubr v1.0.0. Spatial proximity plots were generated by first creating an InSitu dimensional reduction within the Seurat object using centroid coordinates and visually connecting each query cell with its nearest neighbor using plotting features in ggplot2.

For further visualization, 3D UMAPs were also generated using Seurat’s RunUMAP function with the n.components parameter set to 3. The UMAPs were plotted using the Seurat.utils package v2.8.5, with a modified version of the function plot3D.umap. The generated plots were exported to portable HTML webpages and embedded into a webpage developed with the NextJS library (v13.3) using the AppRouter system. The webpage was uploaded to a GitHub repository and hosted using the GitHub Pages system.

### Rarecyte Orion™ Multiplex Immunofluorescence

The Rarecyte Orion^TM^ multiplex immunofluorescence (MxIF) platform, capable of high-plex, high-throughput spatial profiling, was used to analyze human and mouse paraffin embedded (FFPE) tumor tissues and produce a comprehensive visualization of biomarker expression within the tumor microenvironment. Human tumor tissues were formalin fixed (3.7%), paraffin embedded (FFPE) and serially sectioned at a 5 micron thickness. Orion multiplex immunofluorescence was first performed using an 18-marker antibody panel in addition to Hoechst 33342. After deparaffinization and antigen retrieval (EZ-AR^TM^ 2 Elegance RTU, HK547-XAK) using BioGenex EZ-Retriever System, Autofluorescence Quenching was performed using AF quench buffer (4.5% H_2_O_2_ / 24 mM NaOH in PBS) at room temperature first under light box for 60 minutes and then under UV (UltraBright Dual Wavelength 302/365 nm, 21cm x 26cm) for 30 minutes. Slides were then blocked with Image-iT Signal Enhancer (Thermo Fisher Scientific, I36933) for 15 min at room temperature, washed with Surfactant Wash Buffer (0.025% Triton X-100 in PBS) and stained with antibody panel for 3 hours at room temperature. Combinations of the following Rarecyte biomarker antibodies were used for the 18-plex panel: CD4, CD8a, CD11b, CD31, CD163, Granzyme B, KI67, LAG3, PanCK, PD-1, PD-L1, alpha Smooth Muscle Actin (SMA), SOX10, SPP1, TIGIT, TREM2,and Vimentin; and 14-plex panel SPP1, Ki-67, GFAP, SMA, CD44v9, CD8a, CD11b, TREM2, CD163, PD-L1, SOX10, Pan-CK,and Vimentin. After primary antibody incubation, slides were washed with Surfactant Wash Buffer, and nuclei were subsequently counterstained with Hoechst 33342 at a 1:1000 dilution. Slides were mounted using #1.5 thickness coverslips (VWR, 48393-241) and ArgoFluor^TM^ Mounting Medium (Rarecyte, 42-1214-000), cured overnight, and cleaned prior to scanning. Whole slide images were captured at 20X using the Orion and scans were imported into the Artemis Software. The extraction matrix was applied to the raw data and imported into OMERO Plus and PathViewer (Glencoe Software) for visualization.

#### Selection Strategy for Regions of Interest for CosMx SMI and MxIF Analyses H&E Staining and Selection of ROIs

A H&E staining workflow on a serial section adjacent to the experimental slides was performed for tumor tissue. Tissue integrity and specific morphology and histology before scanning was evaluated. In consultation with the pathologist areas indicative of stages of pancreatic disease were identified based on the WHO series on the classification of human tumors in the digestive system. Histopathological diagnosis was based on the WHO Blue Books that pre-sent the gold standard for diagnosis of tumors^29,30^. All tissues presented, including the TMA, were analyzed and ROIs clearly indicated in the Results section. In our experiments we took an “agnostic grid sampling” approach that covered data collected from the biospecimen for both the CosMx SMI and MXIF analyses. Up to 250 ROIs that covered each biospecimen were used to evaluate every possible cell-to-cell spatial proximity and interaction without selection bias with clear nuclear staining and cell segmentation and avoiding areas of tissue folding or loss of integrity.

#### CosMx SMI Analysis

H&E staining with a serial section before beginning the CosMx SMI experiment was used to evaluate tissue block quality and in consultation with the pathologist guided the placement of the fields of view (FOVs). Based on the morphology markers, areas of tissue autofluorescence were avoided to avoid impact cell segmentation.

#### Orion MxIF Analysis

The Orion platform scans the entire tissue slide of up to 8 cm² in a single round of high-plex fluorescent and brightfield imaging that allows us to benchmark biomarker expression (such as CD44v9) onto the same H&E stained slide. The ROIs for the MxIF analysis were based on the H&E staining on the exact same cell and slide whereby the pathologist chose the ROIs based on classical morphological hallmarks that define different stages of disease progression of PDAC^29,30^.

### Image processing, segmentation and analysis using QuPath

Stitching, channel registration, illumination and geometric distortion correction were performed with Artemis software on the Orion platform, and single-cell data analysis was then performed using MCMICRO modules, including UNMICST2 with cell masks that involved 5-pixel dilation of the nucleus mask. Mean and median intensity of each channel and morphological features were quantified for each cell. MxIF image analysis for CD8a T-cell density visualization was performed in QuPath (version 0.6.0-rc3)^31^ on images patient 009, patient 010 and patient R230170. Regions of interest (ROIs) were selected using H&E-stained slides and MxIF (same slide staining) based on patterns of tumor infiltration and inflammation in conjunction with board-certified pathologists. Whole tissue annotation was performed by hand in addition to ROIs by a board-certified pathologist.

#### HALO Spatial Analysis of Rarecyte Orion™ Multiplex Immunofluorescence: Proximity Analysis

Orion™ MxIF images were imported into the HALO image analysis platform (Indica Labs, Albuquerque, NM, USA). Spatial analyses were restricted to prespecified tissue regions of interest (ROIs) within the Orion MxIF images. Areas containing tissue folds, fluorescence artifacts, or inadequate tissue were excluded before analysis. Each marker channel setting was adjusted and optimized. Prior to proximity analysis, the most optimal markers from the Orion MxIF panel were selected to train the HALO AI Custom module to perform nuclear, membrane, and cytoplasmic segmentation within the Orion MxIF images. Segmentation and phenotyping settings were then optimized for each cell type and visually reviewed against the multiplex fluorescence signals to ensure accurate identification of cellular boundaries and marker-positive populations. Next, Proximity Analysis was used to assess spatial proximity between CD44v9-positive CSCs and neighboring cell populations. Target cell populations were assessed within defined distance ranges from CD44v9-positive CSCs. Distances were reported in micrometers. The number and proportion of target cells within each defined proximity range, along with HALO Spatial Plots and markup images, were generated. Quantitative data generated from the HALO proximity analyses were exported and imported into GraphPad Prism (GraphPad Software, San Diego, CA), where the data were used to generate graphs of the spatial relationships.

### Single Cell RNA Sequencing (scRNA seq)

Patient derived PDAC organoids were collected and dissociated into a single-cell suspension using 0.25% Trypsin/EDTA (Thermo Fisher Scientific 25200056). For single-cell transcriptome isolation, ∼15,000 cells per sample were resuspended in sample buffer (BD Biosciences 65000062) then filtered through a 40 microns cell strainer and were loaded into a BD Rhapsody cartridge (BD Biosciences 400000847). Microbead-captured single-cell transcriptomes were utilized to prepare a cDNA library based on whole-transcriptome analysis performed on single cells using the BD Rhapsody system. Briefly, the microbead-captured single-cell transcriptome generated the double-stranded cDNA through the processes of reverse transcription, second-strand synthesis, end preparation, adaptor ligation, and whole-transcriptome amplification (WTA). The BD Rhapsody cDNA Kit (BD Biosciences, 633773) and the BD Rhapsody Targeted mRNA and WTA Amplification Kit (BD Biosciences, 633801) were then used to construct the final cDNA library from double-stranded full-length cDNA. Then, PE150 mode (paired-end with 150-bp reads) on NovaSeq6000 System (Illumina) was used to sequence the library. Further using the whole transcriptome analysis pipeline on the Seven Bridges Genomics platform about 80,000 reads were demultiplexed, trimmed, mapped to the GRCh38 annotation, and quantified. Quality control and downstream analysis were performed with Seurat version 5^21^. Briefly, cells with less than 250 detected genes or less than 500 counts were excluded from the analysis. Cells with more than 30% of mitochondrial or 20% ribosomal reads were excluded. Mitochondrial and ribosomal genes were removed from further analyses. Data normalization was performed using SCTransform v2 normalization method^32^ regressing out mitochondrial percentage. Following normalization, dimensionality reduction was conducted via principal component analysis (PCA), and Uniform Manifold Approximation and Projection (UMAP) was performed for initial visualization. Data was integrated using the IntegrateLayers v5 function with the reciprocal PCA (RPCA) method and SCT normalization. Post-integration, nearest neighbors were identified in the integrated space using the FindNeighbors function and cells were clustered in Seurat using the FindClusters function with the Louvain algorithm (resolution of 0.7). Differentially expressed genes for each cluster were identified using a Wilcoxon rank-sum test and FindAllMarkers function. Cell type annotation was performed using specific gene marker sets and identified cell type markers from published PDAC scRNAseq studies^33^.

### Orthotopically transplanted Nod scid gamma mice using patient-derived organoids

All mouse studies were approved by the University of Arizona Institutional Animal Care and Use Committee (IACUC) Protocol 19-571 that maintains an American Association of Assessment and Accreditation of Laboratory Animal Care (AAALAC) facility. Nod scid gamma mice (The Jackson Laboratory, Stock number 005557) were orthotopically transplanted with approximately 500 patient-derived pancreatic cancer organoids according to our previously published protocols ^14,20^.

### Access to high-resolution spatial transcriptomic data and histology

High-resolution spatial transcriptomic maps and histology can be accessed at https://zavros-lab.github.io/pdac-gallery/. The webpage was developed as a standard HTML-CSS-JavaScript site, using the OpenSeaDragon (v5.0.1) library to render the high-resolution images and provide a zoomable viewer for them. A custom event hook was implemented to synchronize the three image viewers (H&E, spatial plots, image plots) so the viewers displayed the respective data for the same sample. Site assets and code were stored in a GitHub repository, and the webpage was hosted using the GitHub Pages static-site hosting service, making it available to browse using the link mentioned previously.

### Statistical Analysis

Significance of the results was tested using a commercially available software (GraphPad Prism, GraphPad Software, San Diego, CA). Analysis was performed using one-way or two-way ANOVA where a p value of less than 0.05 was determined significant. Sample size for each experiment was determined using a power analysis with SigmaStat software. A power greater than 0.8 was considered effective to determine significance.

## RESULTS

### Spatial profiling reveals distinct cellular niches and persistent cancer stem cells in PDAC tissues

To investigate the cellular composition and spatial organization of the PDAC tumor microenvironment (TME), we applied CosMx**™** Spatial Molecular Imager (SMI) and Rarecyte Orion multiplex immunofluorescence to formalin-fixed, paraffin-embedded (FFPE) tumor tissue sections collected from patients at various stages of disease progression. This workflow enabled high-resolution mapping of cell populations and molecular features within the TME, which we benchmarked against patient-derived organoids (PDO) models (**Fig. 1A-D**).

**Figure 1:**
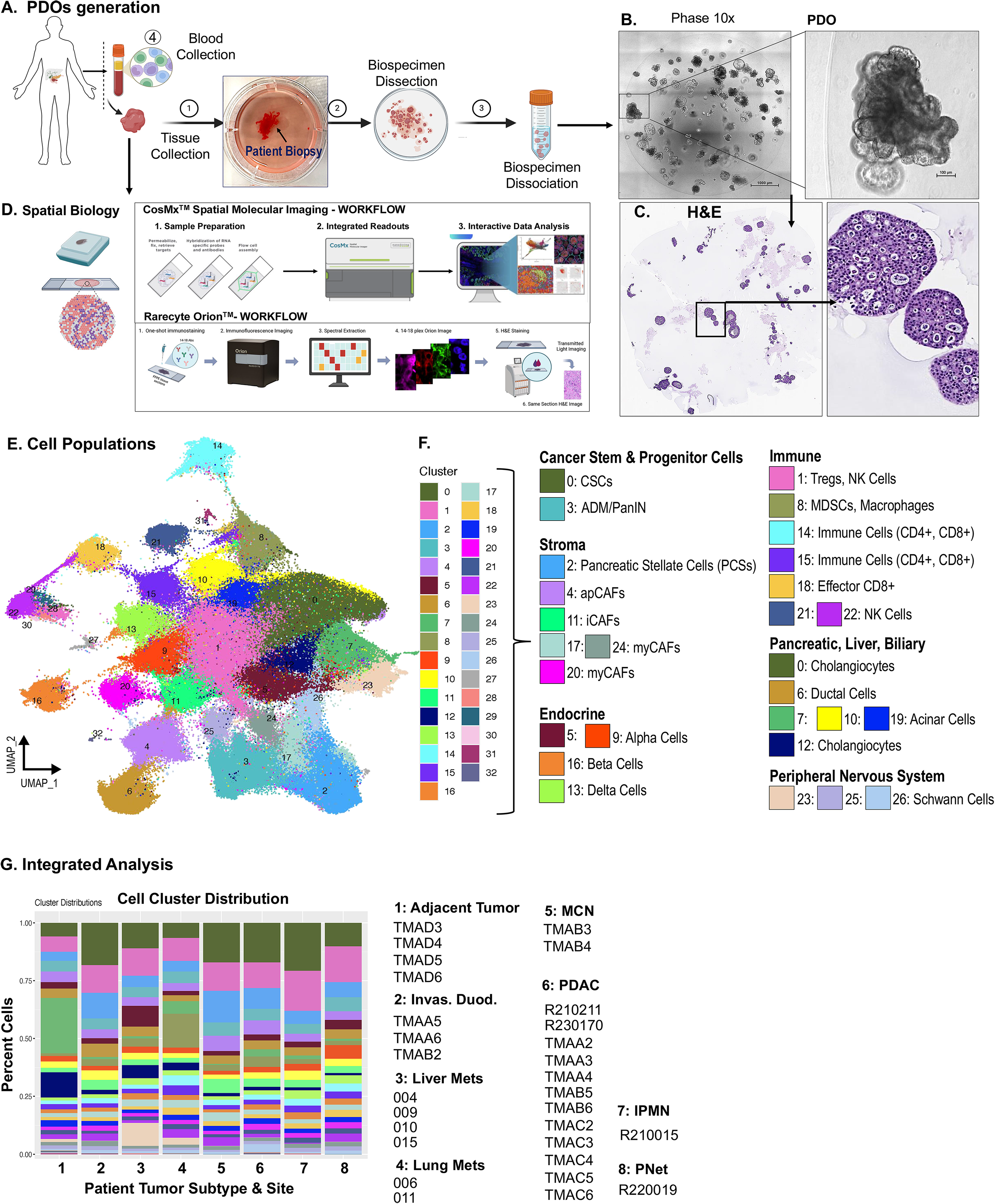
Workflow for the high-resolution mapping of cell populations and molecular features within the TME benchmarked against patient-derived organoids (PDO) models. (A-C) Overview of methodology implemented for the generation of PDO/CAF/IMM cell co-cultures and **(D)** spatial biology. **(E)** Integrated UMAP plot showing cell populations across all patient tumor tissues and fields of view (FOV) selected. **(F)** Identified cell populations based on known gene markers and separated into categories of cancer stem and progenitor cells, stroma, endocrine, immune and pancreatic, liver and biliary specific cell populations. **(G)** Cell distribution across niches 1-7 across all biospecimens analyzed within disease tissues and stages 1) adjacent tumor or cancer (AC), 2) Invasive duodenum, 3) liver metastatic lesions, 4) lung metastatic lesions, 5) mucinous (MCN), 6) PDAC, 7) IPMN and 8) pancreatic neuroendocrine tumor (PNet). Scale bars shown in (B) 1000μm and high magnification image 100 μm.

Unsupervised clustering of gene profiles from spatial transcriptomics identified 33 cell subsets (0-32) (**Fig. 1E, F**). These clusters were annotated using established gene signatures classified the clusters into broad categories of cancer stem and progenitor cells, stroma, endocrine, immune, pancreatic, liver and biliary, and peripheral nervous system (**Fig. 1F**). The top differentially expressed genes were identified for each cell type (**Supplemental Figure 2**) and was used to identify cell population clusters 0 to 32 (**Fig. 1E, F**). The analysis covered 8 disease stages and/or sites that included 1) adjacent tumor, 2) invasive duodenum, 3) liver metastases, 4) lung metastases, 5) mucinous cystic neoplasm (MCN), 6) PDAC, 7) intraductal papillary mucinous neoplasm (IPMN) and 8) one pancreatic neuroendocrine tumor (PNet) (**Fig. 1G**). The cell cluster distribution based on the percentage of cells is shown in **Figure 1G**. Among the pre-dominant cell types that persisted through disease stages were Schwann cells (found predominantly in clusters 23, 25 & 26), cancer stem cells (CSC, cluster 0), immune cells, including Tregs, a diverse phenotype of cancer associated fibroblasts (CAFs) and mesenchymal stem cells (MSCs), pancreatic endocrine cells, MDSCs (cluster 8) and macrophages, hepatic stellate and Kupffer cells, and acinar and ductal cells, of which a subset were identified as undergoing acinar-to-ductal metaplasia (**Fig. 1F, G**). For an interactive 3D UMAP visualization of **Fig 1E-G**, see: https://zavros-lab.github.io/pdac-3d-umap-gallery/.

Spatial data analyzed from patient biospecimens demonstrated a shift in the cell distribution within cellular neighborhoods (niches) in the TME at different stages of disease (**Fig. 2A, B, C**). The network dynamics among precancerous (IPMN), PDAC and metastatic TMEs were analyzed by implementing a network motif analysis (**Fig. 3**). We observed shifting weighted and directed cell circuit interaction networks at different stages of disease (**Fig. 3, Source Files:** https://github.com/zavros-lab/Spatial-Biology-PDAC). These analyses revealed complex interaction networks with CSCs (cluster 0), MDSCs (cluster 8), Schwann cells (clusters 23, 25, 26), and CAFs (clusters 20 myCAFs, 11 iCAFs, 4 apCAFs) as the top sending signals to the other cell types that was dependent on the Niche and stage of disease (**Figs. 3 & 4**). In Niche 1 the strongest interaction occurred among MDSCs receiving signals from CSCs, iCAFs then myCAFs, Schwann cells, pancreatic stellate cells (PSCs), acinar cells and Tregs within the IPMN and liver metastatic (**Fig. 3A**) TMEs. Niche 2, predominantly located in the PDAC and liver metastatic TMEs, the strongest interactions occurred between MDSC and receiving signals from CSCs, myCAFs and PSCs with outgoing signaling from the MDSCs to ADM/PanIN and acinar cells (**Fig. 3C**). The strongest interactions within Niche 3 were between Schwann cells receiving signals from MDSCs, CSCs and acinar cells, in addition to exhibiting autocrine signaling (**Fig. 3E**). Within niches 4 and 5 we identified that the strongest instance of the cell circuit interaction network within the TME of IPMN containing CSCs, myCAFs, iCAFs, MDSCs and T cell lymphocytes (**Fig. 4A**). Circuit analysis of Niche 5 revealed that CSCs supported themselves by means of an autocrine loop, while also signaling to apCAFs and MDSCs, while receiving signals from PSCs, ductal cells and myCAFs (**Fig. 4C**). Within Niche 6 found predominantly in the liver metastatic TME, we found that the strongest interaction occurred between myCAFs and MDSCs, followed by the interaction between myCAFs and Schwann cells and an autocrine loop in which myCAFs, MDSCs and CD8+ lymphocytes potentially send ligands which they also sense by expressing their receptors (**Fig. 4E**). Niche 7 was abundantly located in the TME of IPMN lesions whereby the strongest interaction CSCs was MDSCs, Schwann cells, iCAFs and acinar cells (**Fig. 4G**). CellChat was used to identify the intercellular signaling pathways as an indicator of the functional and regulatory roles of key cell subpopulations (**Source File:** https://github.com/zavros-lab/Spatial-Biology-PDAC). Overall signaling patterns were represented in heatmaps of the pathways from the signals originating from each cell cluster (**Fig. 3B, D, F and Figure 4B, D, F**). The CSC cell population (cluster 0) consistently expressed signaling pathways related to MIF, CEACAM and VEGF (**Figs. 3 & 4**). All source files can be located at this link: https://github.com/zavros-lab/Spatial-Biology-PDAC.

**Figure 2:**
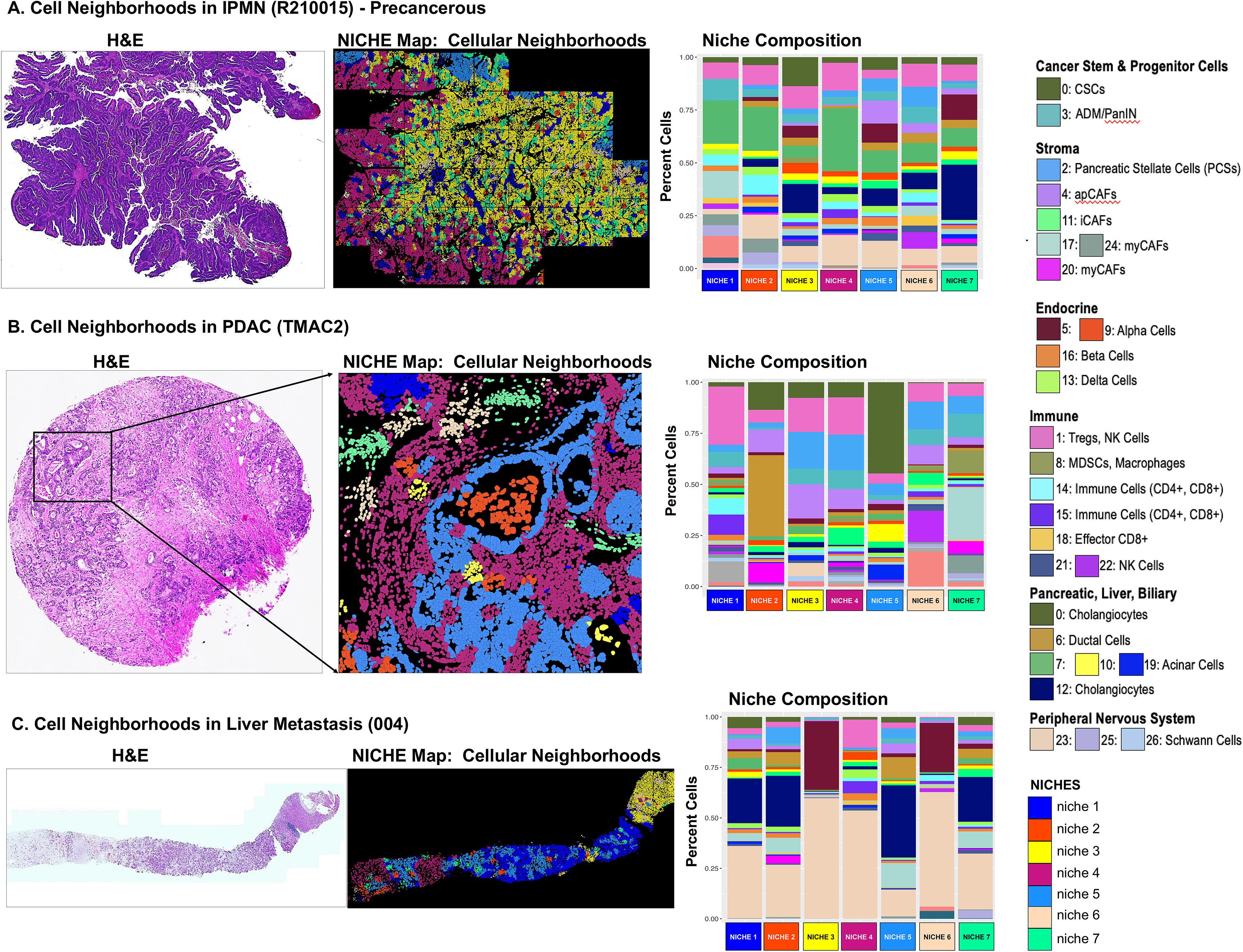
CosMx*™* SMI analysis of the TME at different stages of disease. H&E stain, niche map of cellular neighborhoods and cell distributions across niches 1-7 of **(A)** a precancerous IPMN lesion, **(B)** PDAC, and **(C)** core biospsy collected from a liver metastatic site.

**Figure 3:**
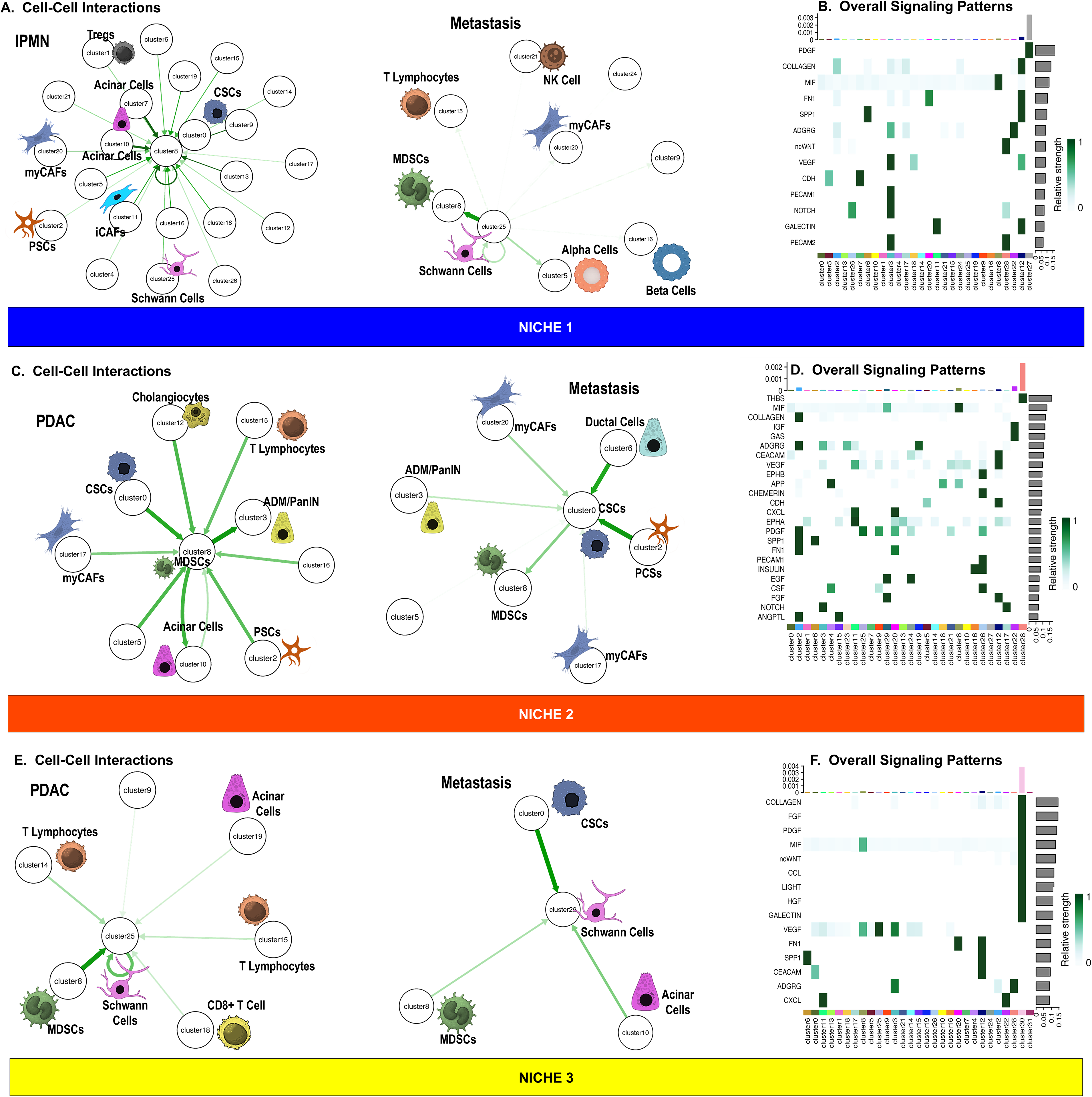
Network motif and CellChat analyses. Weighted and directed cell circuit interaction networks and heatmaps of overall signaling patterns predicted in **(A, B)** niche 1, **(C, D)** niche 2, and **(E, F)** niche 3.

**Figure 4:**
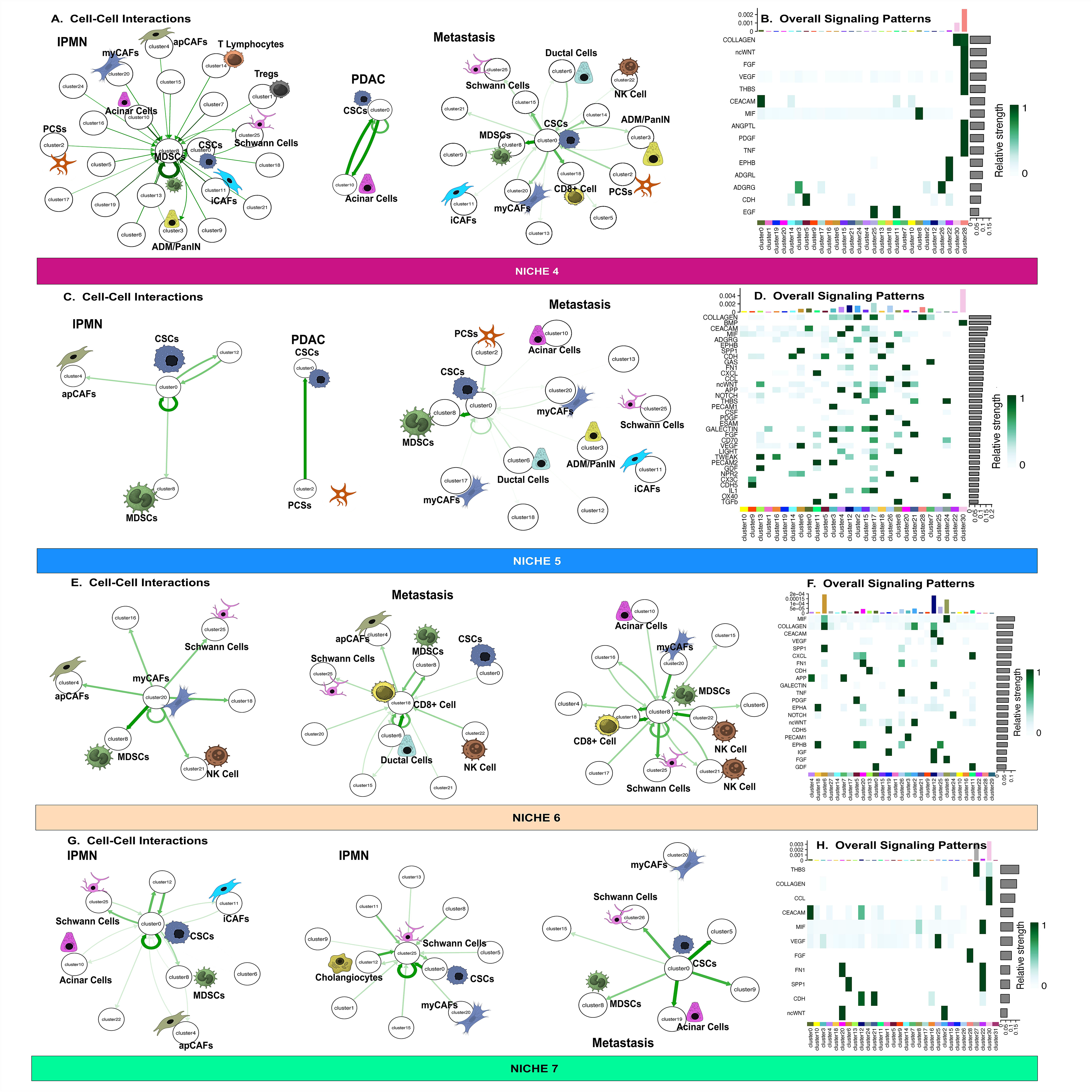
Network motif and CellChat analyses. Weighted and directed cell circuit interaction networks and heatmaps of overall signaling patterns predicted in **(A, B)** niche 4, **(C, D)** niche 5, **(E, F)** niche 6, and **(G, H)** niche 7.

Cell populations identified in **Fig. 1E & F** were spatially mapped across all samples (https://zavros-lab.github.io/pdac-gallery/). Across all tissue sections analyzed, we observed heterogeneity among the cellular neighborhoods (niches) comprising of distinct cell distributions (**Fig. 5A**). Reference to the pathology defined by H&E stain of a tissue microarray (TMA) is shown as a representative example **in Figure 5B**. The cell distributions (**Fig. 5D, G**) within niches (**Fig. 5C, F**) were consistent with the spatial expression of key gene markers shown in feature plots (**Fig. 5E**). Notably, expression of cancer stem cell markers CD44, and TROP2 was observed across all samples, with high expression in areas of cancer cell localization (**Fig. 5E**, niches 5 & 7). Network motif analysis showed that the strongest interactions within niche 5 were between PSC sending signals to myCAFs, apCAFs, ductal cells and CSCs (**Fig. 5H**), and niche 7 with a cell circuit interaction network between CSCs receiving signals from myCAFs and PSCs while sending signals to MDSCs (**Fig. 5I**) between tumor cells and stroma (**Fig. 5J**). NOTCH, Wnt and ADGRG signaling patterns were identified as sending signals originating from the PSCs spatially mapped within niche 5 (**Fig. 4D**).

**Figure 5:**
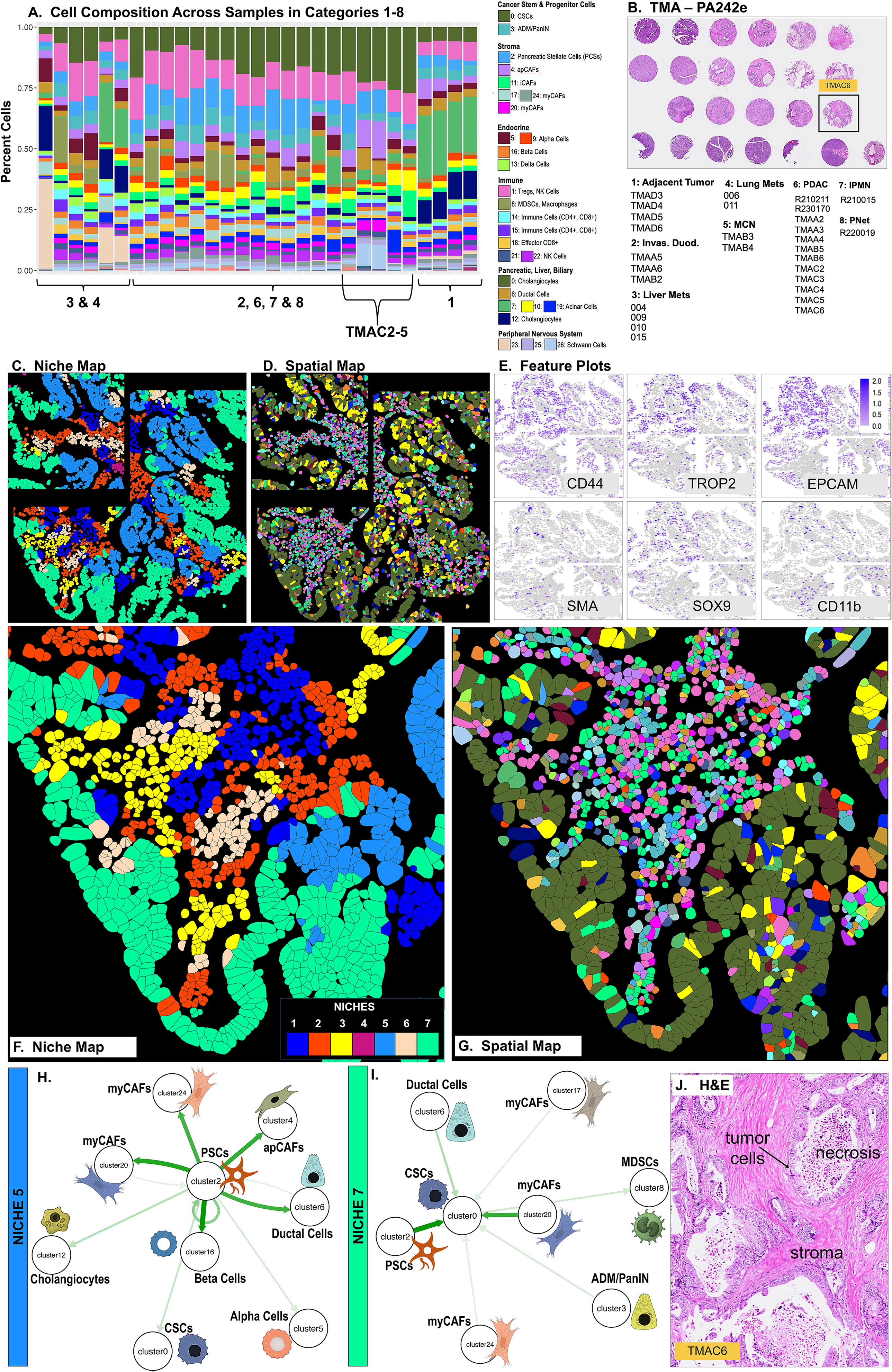
CosMx*™* SMI analyses of surgical resected biospecimens collected from patients with PDAC. (A, **B)** Cell distribution across all tissues at disease stages 1-8 that were analyzed. **(C)** Representative FOVs were selected from PDAC biospecimen TMAC6 and shown as a **(C)** niche and **(D)** spatial maps. **(E)** Feature plots highlight the expression of genes that included alpha smooth muscle actin (SMA), CD44, CD11b, TROP2, EPCAM and SOX9. Higher magnifications of **(F)** niche and **(G)** spatial maps. Weighted and directed cell circuit interaction networks predicted in **(H)** niche 5, and **(I)** niche 7. **(J)** H&E stain of TMAC6.

We next defined the cellular niches and their cell distribution within the TMEs of liver metastatic sites (**Fig. 6A**, patients 009 and 010). The TME of liver metastatic lesions exhibited cell distributions comprising of MDSCs, myCAFs and CSCs (**Fig. 6B-D**). The cell distributions were consistent with the spatial expression of key gene markers shown in the feature plots for cancer stem cell markers CD44, and TROP2, with high expression in areas of cancer cell location (**Fig. 6E**). SMA and vimentin were expressed within the stroma, supporting a predominant myCAF infiltration (**Fig. 6E**). Together, these data demonstrate that spatial profiling of PDAC tissues across disease stages and metastatic sites reveals dynamic, heterogeneous cellular niches, with persistent populations of Schwann cells, CAFs, and CSCs-including those marked by CD44 and TROP2-that may contribute to disease progression and therapeutic resistance.

**Figure 6:**
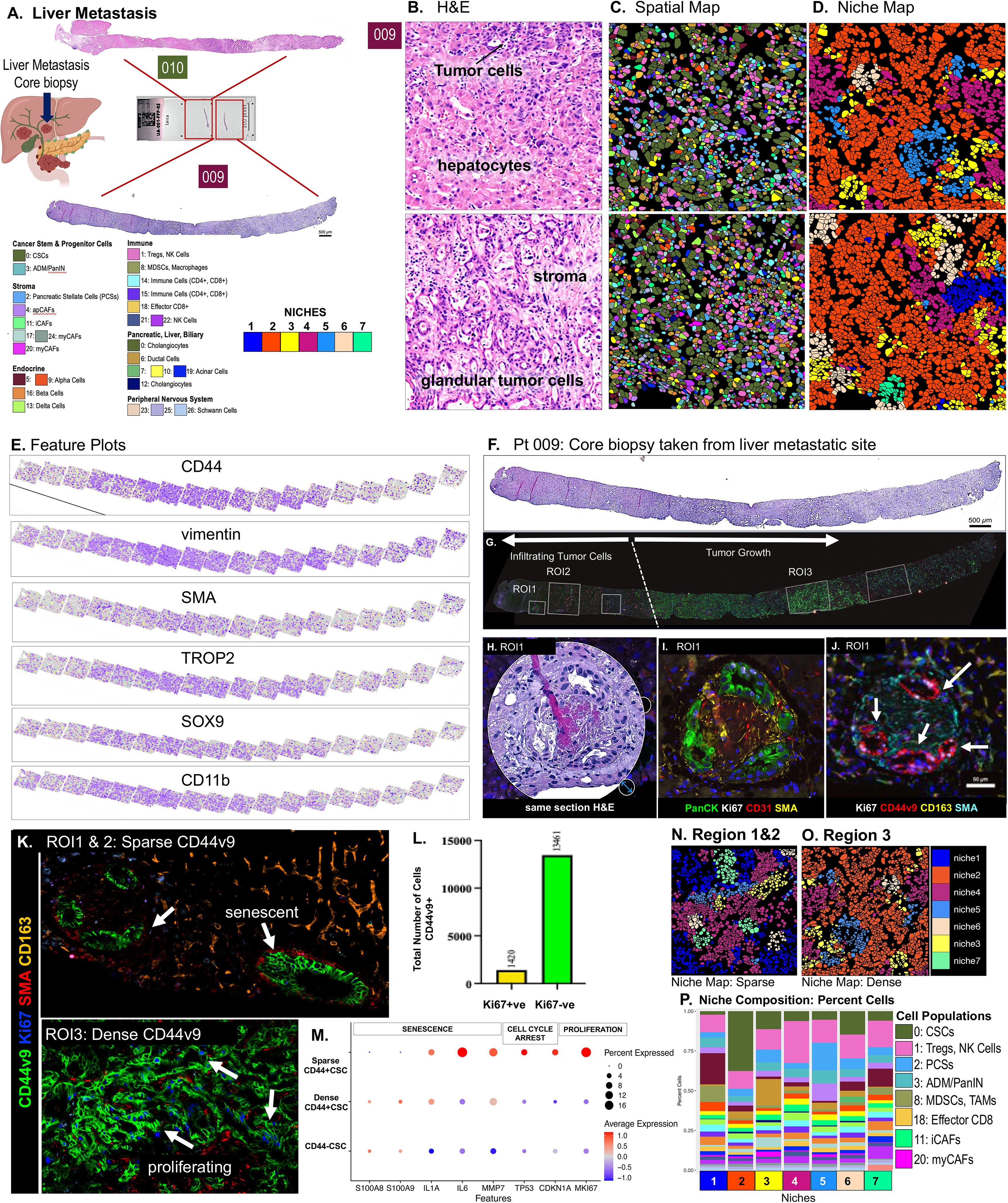
CosMx*™* SMI analyses of core biopsies collected from liver metastatic lesions. (A, **B)** H&E stain of the core biopsies 009 and 010 used in the analyses. Representative FOVs were selected and shown as a **(C)** spatial and **(D)** niche maps. **(E)** Feature plots show highly expressed genes that included SMA, vimentin, CD44, CD11b, TROP2 and SOX9. **(F)** Scan of H&E-stained slide of patient core biopsy 009 from a liver metastatic site. **(G)** Highlighted ROIs 1-3 based on distinct cell types and tumor stromal features on MxIF image. **(H)** Same H&E-stained section highlighting ROI1. MxIF staining highlighting the expression of **(I)** PanCK (green), CD31 (red), SMA (yellow), Hoechst (blue), and Ki67 (white). **(J)** Expression of CD44v9 (red) in the same cancer cells in ROI1. Scale bars shown in (F, G) 500μm and (J) 50 μm. **(K)** ROIs 1-3 shown at higher magnification whereby arrows indicate Ki67 negative senescent or Ki67 positive proliferating CD44v9 positive cells with quantification shown in **L**. **(M)** Dot plot of the average expression of markers of senescence, cell cycle arrest and proliferation in sparse and dense CD44v9 CSCs. Representative FOVs were selected and shown as a niche maps from **(N)** region 1&2, or **(O)** region 3. **(P)** Percent cell distribution across niches.

#### CD44v9 is expressed on a cancer stem cell population identified in post-SOC surgical biospecimens and persists at sites of liver metastases

The spatial biomarkers identified by SMI were validated and benchmarked to the patient’s tumor pathology using a designed Orion antibody panel. A FFPE tumor tissue section was prepared from a core biopsy collected from patient 009 diagnosed with liver metastatic PDAC (**Fig. 6F**) and ROIs were selected (**Fig. 6G**). Expression of CD44v9 (red) positive cells were localized specifically to the tumor regions (**Fig. 6H-J**). In ROIs 1 and 2 CD44v9 positive cells were localized within the liver tissue in “nests” and described as “sparse” (**Fig. 6G, K**). In ROI 3, CD44v9 positive cells were described as “dense” tumor regions (**Fig. 6G, K**). We observed that there were two populations of CD44v9 cells that were either Ki67 positive proliferating cells, or a population that was morphologically senescent and stained negative for Ki67 (**Fig. 6J, K and indicated with arrows**). Quantification of fluorescence intensity using HALO, showed that the number of CD44v9 positive cells that were non-proliferating and negative for Ki67, were significantly greater across the tissue (**Fig. 6L**). Analysis of genes expressed in senescent cells and indicative of cell cycle arrest were significantly elevated in CD44+CSCs within the sparse region compared to those cells expressed in the dense region (**Fig. 6M**). Interestingly, proliferative marker MKi67 was significantly increased in the sparse region (**Fig. 6M**). Given that the dot plot calculates average expression relative to the other comparison, and since there are more CD44+ CSCs in the dense region that don’t express MKI67, the overall average was lowered (**Fig. 6M**). While the TME of liver metastatic lesions exhibited cell distributions comprising of MDSCs, myCAFs and CSCs (**Fig. 6B-D**), a closer analysis of these regions showed distinct cell niches comprising of a shift in the distribution of the cellular neighborhoods (**Fig. 6N, O, P**). In the area of sparsely CD44v9 sparsely distributed (region/RO11&2), one of the predominant cellular neighborhoods observed was niches, 1,3 and 4 comprising of predominantly Tregs, NK cells, immunosuppressive MDSCs and TAMs (**Fig. 6N, P**). In comparison, the area of densely distributed CD44v9 populations (region/ROI3), the predominant cellular neighborhood observed was niche 2 comprising of an expansion of the CSCs population (**Fig. 6O, P**).

Network motif analysis was implemented to identify differences in the TME surrounding CD44v9 proliferating and non-proliferating positive cells whereby there was a shift in the weighted and directed cell circuit interaction networks within the regions of the sparsely and densely expressed CD44v9 CSC populations (**Fig. 7A, B, C, D**). In the region of sparse CD44v9 non-proliferating cells, there were significant outgoing signals from the CD44 CSC population (cluster 0) to MDSCs, effector CD8 CTLs, and myCAFs (**Fig. 7C**). Within the region of dense proliferating CD44v9 cells, CD44 CSCs predominantly received signals from myCAFs, PSCs, and ADM/PanIN lineage cells, with outgoing signals to MDSCs (**Fig. 7D**). Nearest neighbor proximity analysis of the transcriptomic data (**Fig. 7B**) showed that, compared to the region of sparsely expressed CSC population (cluster 0) (**Fig. 7E**), significantly greater numbers of the CD11b+ MDSCs and CD8+ cells and myCAF and iCAFs were found within 250 μm of the CD44+ CSCs located in the dense region (**Fig. 7F**). Further CellChat signaling analysis showed that the outgoing signaling pathways from the sparse CD44+ CSC population included chemokine CCL, glycoprotein CSF, and cytokine MIF (**Fig. 7G**). In the dense CD44+ CSC region, predominant receiving signals included fibronectin, SSP1, and collagen (**Fig. 7H**).

**Figure 7:**
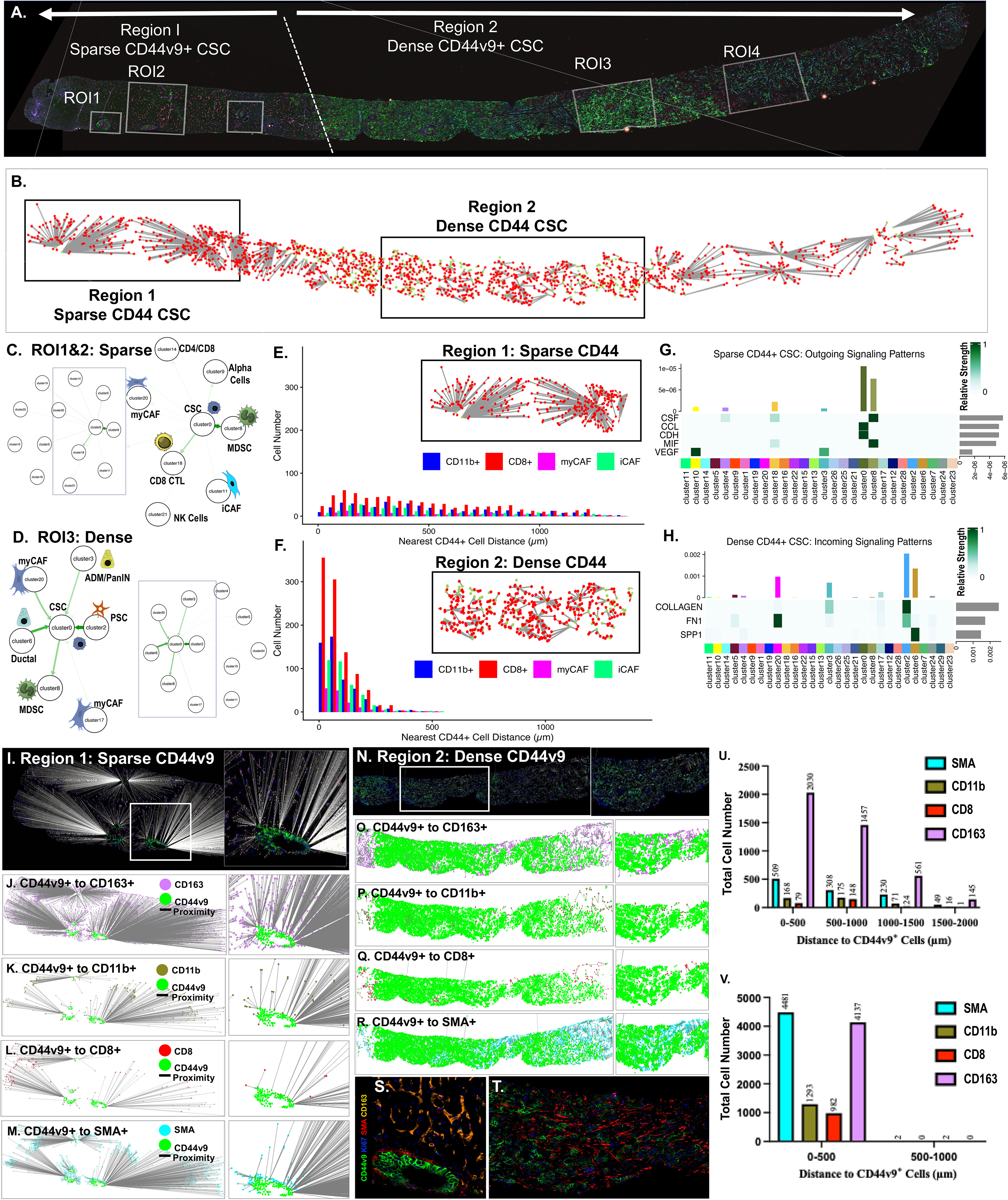
Proximity analyses of transcriptomic and MxIF data. Images of patient 009 tissue used for proximity analyses of **(A)** MxIF data, and **(B)** CosMx transcriptomic data. Network motif and CellChat analyses of weighted and directed cell circuit interaction networks of overall signaling patterns predicted in **(C)** ROI1&2 and **(D)** ROI3. Proximity analysis of transcriptomic data across **(E)** ROI1&2 and **(F)** ROI3 measuring distance of CD11b, CD8 myCAFs and iCAFs from CD44+ CSCs (cluster 0). Heatmaps of overall signaling patterns predicted as **(G)** outgoing from sparse CD44+ CSCs and **(H)** incoming from dense CD44+ CSCs. HALO proximity analysis measuring distance of CD11b, CD8 myCAFs and iCAFs from CD44v9 CSCs in **(I-M)** sparse region 1 (ROIs1&2) and **(N-R)** dense region 2 (RO13). Quantification of proximity analyses in **(U)** sparse region 1 (ROIs1&2) and **(V)** dense region 2 (RO13).

Using the HALO Spatial Analysis Module the proximity from CD44v9+ CSCs (cluster 0) and target neighboring cells that included CD8+ CTLs, CD11b+ MDSCs, CD163+ macrophages and α-SMA+ fibroblasts (myCAFs) was calculated in patient core biopsy 009 collected from a liver metastatic region 1 (sparse, **Fig. 7A, I-M**) and region 2 (dense, **Fig. 7A, N-R**). There were differences in the spatial distance between CD8+ and CD11b+ cells to CD44v9 within the cell niches identified in ROI1&2 (“sparse CD44v9 positive cells) compared to ROI3&4 (“dense” CD44v9 positive cells) (**Fig. 7**). In support of the interaction of myCAFs with CD44v9+ (**Fig. 7S, T**), the significant number of α-SMA+ myCAFs were measured to be within 500 μm of the CSC population (**Fig. 7U, V**). Overall, within the cell number within 500 μm of the CD44v9 positive CSC population was significantly greater in the dense CD44v9 CSC population (**Fig. 7V**), compared to the sparse region (**Fig. 7U**). Spatial proximity of CD44v9⁺ cells to immune and stromal cell populations was also performed in using tissue collected from a liver metastatic lesion of a patient with PDAC. Data was consistent with the observations made with patient 009 (**Supplemental Figure 7**).

Further analysis of the surrounding tissues showed that CD44v9 was absent from the surrounding liver tissue as shown as marked by distinct areas of hepatocytes and Kupffer cells (yellow) (**Fig. 8A, B**). Within ROI1(**Fig. 8C-E**) and ROI3 (**Fig. 8F, G**), GFAP positive cells (magenta) were intricately associated with CD44v9 positive cancer stem cells (green) that was consistent with the detection of Schwann cells by SMI. Network motif analysis within ROI1a (**Fig. 8H**) and ROI3 (**Fig. 8I**), of the complex network of interactions within the TME, within niche 3 a two-cell circuit communication between CD44v9 positive CSCs and GFAP positive Schwann cells participated as one of the highest scoring interaction instances (**Fig. 8J**). Using the “CellChat” package we predicted that CSCs (cluster 0) significantly communicated with the Schwann cells (cluster 25) through the MIF and CEACAM signaling pathways (heatmap shown in **Fig. 3F**, **Fig. 8J**). Further network motif analysis of the niches within the ROI1a (**Fig. 8H**) and ROI3 (**Fig. 8I**) revealed that among the strongest interactions within niche 2 were between myCAFs and CSCs, NK cells and CD8+ lymphocytes (**Fig. 8J**). In this instance, myCAFs (cell cluster 20) within niche 2 significantly communicated with the surrounding cell populations (CSCs, NK cells, CD8+ lymphocytes) predominantly through the FN1 (fibronectin), PDGF, CXCL and EPHA (Ntn1, netrin 1) signaling pathways (heatmap shown in **Fig. 3D**, **Fig. 8J**).

**Figure 8:**
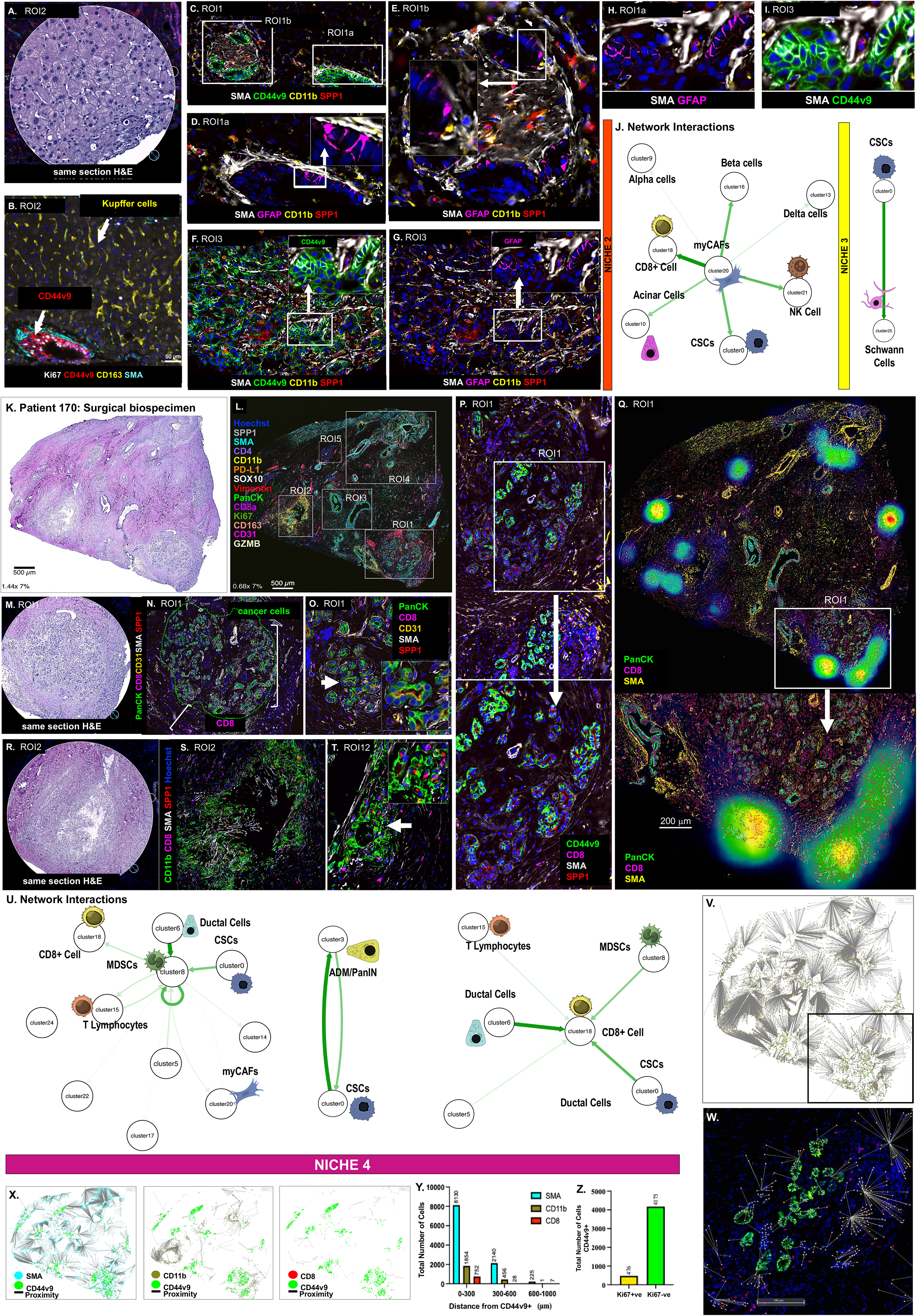
Expression of cancer stem and immune cell markers in a core biopsy collected from a liver metastatic sites and surgically resected tissue biospecimen using the Orion ^TM^ MxIF imaging. **(A)** H&E-stained section of ROI2 from patient 009 showing the expression of CD44v9 (red), Ki67 (while) and SMA (cyan) in **B. (C)** Selected ROI1a and ROI1b with CD44v9 (green) expressing cancer cells. **(D)** Expression of GFAP (magenta) positive cells in proximity of cancer cells in ROI1a (arrow indicates region shown in inset). **(E)** Expression of GFAP (magenta) positive cells in proximity of cancer cells in ROI1b (arrow indicates region shown in inset). **(F)** CD44v9 (green) expressing cancer cells and GFAP (magenta) positive cells within ROI3 (arrow indicates region shown in inset). SPP1 (red) expressed within CD11b infiltrating MDSCs. **(G)** Expression of GFAP (magenta) positive cells in proximity of CD44v9 ex-pressing cancer stem cells in ROI3 (arrow indicates region shown in inset). Higher magnification of ROIa and ROI3 are shown in **H & I** respectively**. (J)** Network interactions predicted in niches 2 and 3. **(K)** Scanned H&E-stained slide of patient surgical biospecimen 170. **(L)** ROIs 1-5 highlighting distinct cell types and tumor stromal features using MxIF. **(M)** H&E section of ROI1. MxIF staining highlighting the expression of **(N)** cancer cells (PanCK, green) surrounded by CD8 cells (magenta). **(O)** Higher magnification of ROI1 showing expression of SPP1 (red) within cancer cells (arrow indicates region shown in inset). **(P)** MxIF staining of ROI1 showing a population of PanCK+/SPP1+ tumor cells co-expressing CD44v9. Higher magnification of area shown in **P** is represented in the inset (arrow indicates region shown in inset). **(Q)** CD8+ ‘hotspots’ within the TME. **(R)** H&E section of ROI2. **(S)** MxIF staining showing the expression of SPPI (red) expression in CD11b (green) positive MDSCs. Higher magnification of area shown **S** is represented in **T** (arrow indicates region shown in inset). **(U)** Network interactions predicted in niche 4. **(V)** Representative image of proximity analysis of patient 170 overlaid with CD44v9 higher magnification shown in panel **W. (X)** HALO proximity analysis measuring distance of CD11b, CD8, and SMA from CD44v9 CSCs. **(Y)** Quantification of proximity analyses. **(Z)** Quantification of Ki67 negative senescent or Ki67 positive proliferating CD44v9 positive cells.

Another representative FFPE tumor tissue section, collected from patient R230170 (170) during Whipple resection, was used to select Regions of Interest (ROI) (**Fig. 8L**) based on histologically relevant areas of the same section H&E post-multiplex immunofluorescence (MxIF) (**Fig. 8K**). Within ROI1 we observed PanCK positive tumor cells (green) that expressed SPP1 (red) embedded within a dense SMA positive stromal compartment (white) (**Fig. 8M-O**). Further analysis revealed, a population of PanCK+/SPP1+ tumor cells co-expressed cancer stem cell marker CD44v9 (**Fig. 8P**). Surrounding the presence of the dense desmoplastic stroma were CD8 positive cells (magenta) (**Fig. 8Q**). CD8+ T cells were segmented using QuPath image analysis to generate a density map showing ‘hotspots’ of CD8+ T cell clusters located in the TME surrounding the tumor cells (**Fig. 8Q**). SPP1 was also co-expressed within CD11b positive myeloid cells within ROI2 (**Fig. 8R-T**). Network motif analysis showed that within niche 4 among the strongest interactions observed included a complex communication network between MDSCs (cluster 8) and myCAFs, T lymphocytes (CD8+ & CD4+), CSCs and ductal cells (**Fig. 8U**). A two-cell circuit communication between CSCs (cluster 0) and a cell population consistent with an ADM/PanIN (cluster 3) gene expression signature was a bidirectional interaction (**Fig. 8U**). Incoming signals to the CSCs from ADM/PanIN cells were predominantly mediated by adhesion G-protein coupled receptor (ADGRG), MIF and CEACAM signaling pathways (heatmap shown in **Fig. 4B**, **Fig. 8J**). Within niche 4, CD8+ lymphocytes (cluster 18) received signaling from surrounding CSCs and MDSCs mediated through incoming predominantly mediated by MIF and CEACAM signaling pathways (heatmap shown in **Fig. 4B**, **Fig. 8J**). Using the HALO Spatial Analysis Module the proximity from CD44v9+ CSCs (cluster 0) and target neighboring cells that included CD8+ CTLs, CD11b+ MDSCs, and a-SMA+ fibroblasts (myCAFs) was calculated in patient biospecimen 170 (**Fig. 8V, W**). The total number of cells was greatest within 500 mm of the CD44v9 positive CSC population (**Fig. 8X, Y**). Quantification of fluorescence intensity using HALO, showed that the number of CD44v9 positive cells that were non-proliferating and negative for Ki67, were significantly greater across patient tissue 170, similar to that observed in patient biospecimen 009 (**Fig. 8Z**).

### A tumor cell population expressing CD44v9 is present within the tissue collected adjacent to tumor

SMI data revealed predominantly acinar (clusters 7, 10 & 19) and ductal (cluster 6) cells, with persistent cells with ADM gene signature (cluster 3), iCAFs (cluster 11) and CSCs (cluster 0) within the tissues adjacent to tumor/cancer (**Fig. 9A-E**). The cell distributions within the TME were consistent with the spatial expression of gene markers shown in the feature plots for cancer stem cell markers CD44, TROP2 and SOX9 (**Fig. 9F**). To investigate whether acinar-to-ductal metaplasia (ADM), a precursor lesion in PDAC, contributes to early neoplastic transformation, we analyzed regions of interest (ROI) in adjacent “normal” tissues. Further analysis of the SMI data showed the co-expression of CD44 and ductal marker CK19 (KRT19) in the same cell populations identified as ADM lesions (cluster 7, 10 & 19) (**Fig. 9G**). Significantly higher expression of CK19 and CD44 was measured in clusters 6 and 0, that were identified as ductal (cancer stem cells) and cancer stem cells, respectively (**Fig. 9G**). Immunofluorescence staining in ROI1a identified cells co-expressing the acinar cell marker amylase and cancer stem cell (CSC) marker CD44v9, alongside CD44v9+/CK19+ cells lacking amylase (**Fig. 9H, I**). A similar population was observed in ROI1b (**Fig. 9J**). In serial sections, cells co-expressing amylase and the ductal marker CK19 were also positive for CD44v9 (**Fig. 9K, L, ROI2**), consistent with metaplastic transition. The FFPE tissue collected from adjacent tumor that was matched from patient R230170 (170) during Whipple re-section, was used to select ROIs based on histologically relevant areas representing tumor or PanIN lesions (**Fig. 9M, N**). Within ROI1a & b we observed CD44v9 positive cells (green) that were embedded within a dense SMA positive stromal compartment (red) (**Fig. 9O-R**). Surrounding the presence of the dense desmoplastic stroma were CD8 (magenta), CD11b (yellow) and CD163 (cyan) positive immune cells (**Fig. 9O-R**).

**Figure 9:**
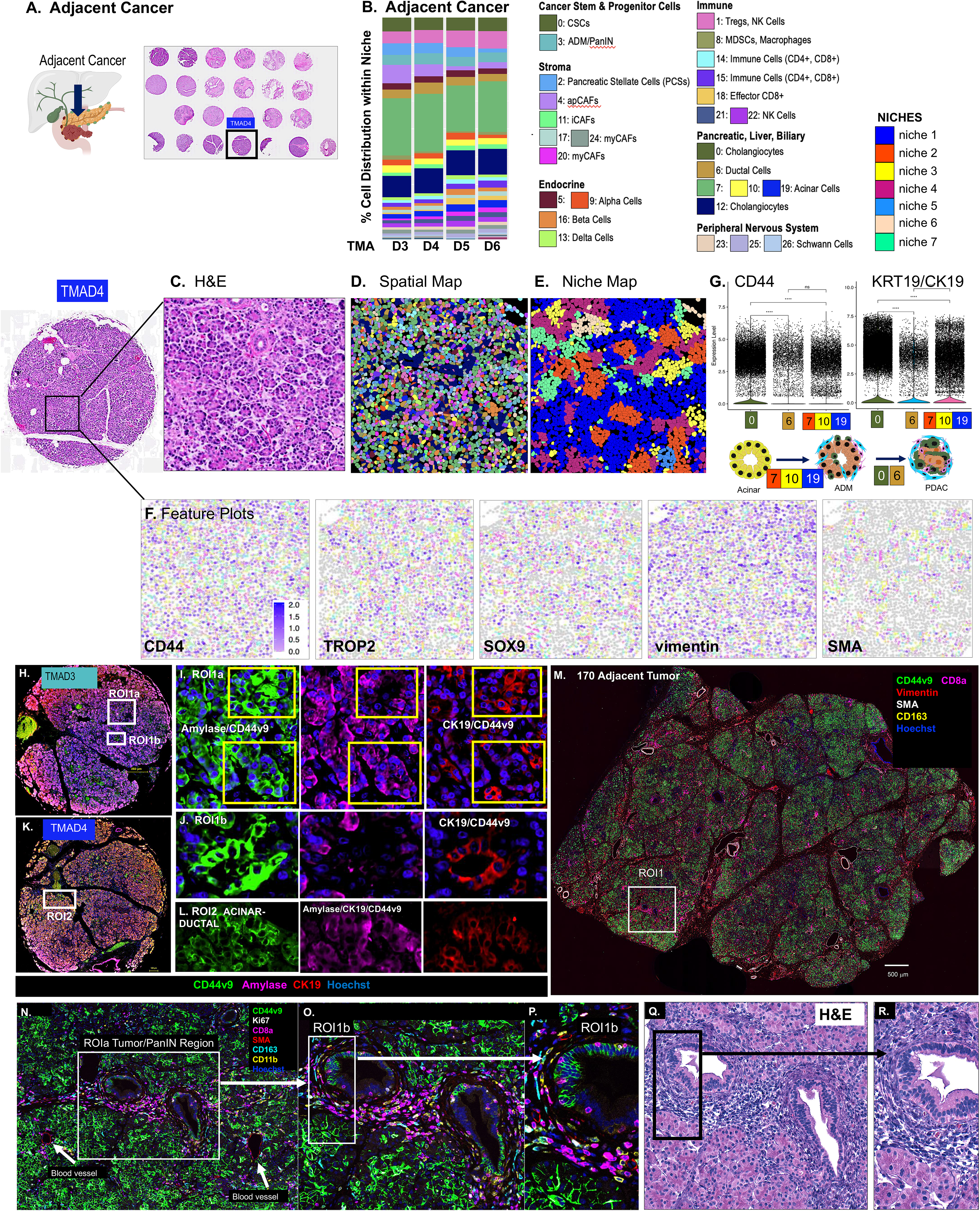
CosMx*™* SMI analyses using surgically resected biospecimens collected from tissue adjacent to the cancer/tumor of patients with PDAC. (A) H&E stain of the TMA used in the analysis. **(B)** Cell distribution across all biospecimens collected from tissue adjacent to the tumor that were analyzed. Representative FOVs are shown for **(C)** H&E stain, **(D)** spatial and **(E)** niche maps. **(F)** Feature plots highlight the expression of genes that included SMA, vimentin, CD44, TROP2 and SOX9. (H, K) Selected ROI from the confocal slide scanned images of patient TMAD3. (G) Gene expression levels of CD44 and CK19 (KRT19) within or across clusters 0, 6 and 1, 10 and 19. Schematic diagram highlighting the expression of CD44v9 in ADM lesions is shown in **G**. Immunofluorescence staining using antibodies specific for detecting the expression of CD44v9 (green), amylase (magenta) and cytokeratin 19 (CK19, red) in corresponding **(H)** ROI1a and ROI1b, **(I)** higher magnification of ROI1a, **(J)** higher magnification of ROIb **(K)** ROI2, **(L)** high magnification of ROI2. **(M)** Scanned MxIF staining of tissue collected from adjacent tumor of surgical biospecimen 170. **(R)** MxIF staining highlighting the expression of CD44v9 (green), Ki67 (white), CD8a (magenta), SMA (red), CD163 (cyan), CD11b (yellow) and Hoechst (blue) in ROIa. Higher magnification of ROI1 is shown in **N**. Higher magnification of ROIa is shown in **O & P**. **(Q & R)** H&E stain of ROI1b at low and high magnification. Scale bar shown in (M) 500μm.

Our data suggest that CD44v9+ CSCs may arise in pre-malignant ADM lesions adjacent to tumors and within IPMNs. The data also raised the question of what renders this CSC population resistant. Therefore, CosMx whole transcriptomic (WTx) SMI approach was taken to further define the CD44v9 positive CSC population as a potential reservoir for tumor recurrence (**Fig. 10**). Unsupervised clustering of gene profiles from spatial transcriptomics identified 28 cell subsets (0-27) (**Fig. 10A**). These clusters were annotated based on established gene signatures classified the clusters into broad categories of cancer stem and progenitor cells, stroma, endocrine, immune, pancreatic, liver and biliary, and peripheral nervous system (**Fig. 10A**). The top differentially expressed genes were identified for each cell type (**Supplementary Figure 2**) and was used to identify cell population clusters 0 to 27 (**Fig. 10A**). The analysis covered 4 disease stages that included 1) PDAC, 2) intraductal papillary mucinous neoplasm (IPMN), 3) mucinous cystic neoplasm (MCN), and 4) cancer adjacent pancreas tissue (AC) (**Fig. 10B**). The cell cluster distribution based on the percentage of cells showed that among the predominant cell types within the TME of AC, IPMN, MCN and PDAC included CSCs, niches comprised of ductal/stroma/and iCAFs, mesenchymal stem cells (MSCs) and Schwann cells (SCs) that were found in the same cellular neighborhood as iCAFs and ductal cells (**Fig. 10C, D**).

**Figure 10:**
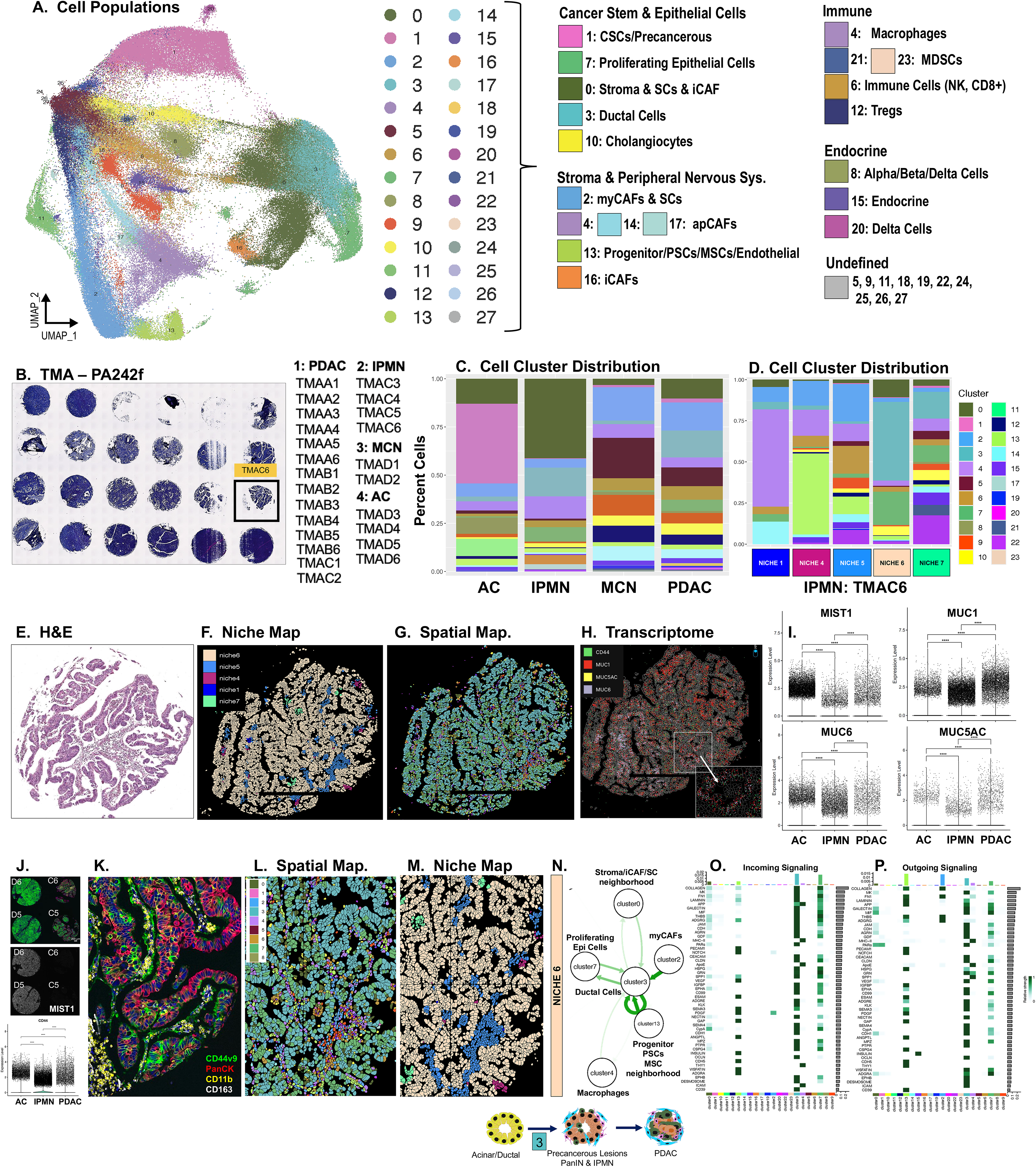
CD44v9 positive CSC population expression in early precancerous lesions. (A, **B)** Integrated UMAP plot showing cell populations across all fields of view (FOV) selected from TMA biospecimens. Identified cell populations based on known gene markers and separated into categories of cancer stem and progenitor cells, stroma and peripheral nervous system, immune, endocrine and undefined populations. **(C)** Cell distribution across all biospecimens analyzed within disease tissues and stages adjacent cancer (AC), IPMN, MCN, and PDAC. **(D)** Cell cluster distribution across niches 1-7 in biospecimen TMAC6 IPMN. Representative FOV were selected and shown in **(E)** matched H&E stain, **(F)** niche and **(G)** spatial maps. **(H)** Spatially mapped transcriptome expression for CD44 (green), MUC1 (red), MUC5AC (yellow) and MUC6 (grey). **(I)** Spatially mapped transcriptome expression for MIST1, MUC1, MUC5AC and MUC6 were quantified. **(J)** Scanned MxIF staining of tissue collected from adjacent tumor and IPMN precancer tissues and quantification of CD44 transcriptome expression in AC, IPMN and PDAC tissues. **(K)** MxIF staining highlighting the expression of CD44v9 (green), CD163 (white), PanCK (red), CD11b (yellow) and Hoechst (blue) in C6. FOV shown in **K** represented as matched **(L)** spatial and **(M)** niche maps. **(N)** Weighted and directed cell circuit interaction networks and **(O, P)** heatmaps of outgoing and incoming signaling patterns predicted in cell clusters within niche 6. Schematic diagram highlighting the expression of cluster 3 in ADM lesions is shown in **N**.

Cell populations identified in **Fig. 10A** were spatially mapped across all samples and observed heterogeneity among niches comprising of distinct cell distributions (**Fig. 10E-G**). Reference to the pathology, shown is a representative example of IPMN and PanIN precancerous lesions (TMAC6) (**Fig. 10E-G**). Mucin expression patterns are often followed as a criterion for classification of precancerous lesions including PanINs and IPMN ^34^. In addition, it is reported that loss of transcription factor MIST 1 leads to enhanced PanIN development, whereby ADM is a highly plastic process involving transdifferentiation and reduced expression of acinar-specific genes including MIST1 ^35^. Therefore, we spatially mapped the expression of MUC1, 5AC and 6, and MIST1 with total CD44 in areas of IPMN lesions (**Fig. 10H**). Overall quantification of transcripts revealed that MIST1 expression was significantly higher in AC tissues compared to IPMN lesions and PDAC (**Fig. 10I**). While expression of MUC5AC was not significantly different among the 3 stages of disease, MUC1 and MUC6 expression was lower in IPMN lesions compared to AC and PDAC (**Fig. 10I**). CD44 expression was significantly higher in PDAC compared to AC and IPMN tissues (**Fig. 10J**). Notably, expression of cancer stem cell marker CD44v9 was observed in areas of MUC1 and MUC6 expression (**Fig. 10J-M**). Network motif analysis showed that the strongest interactions within niche 6 was between cluster 3, identified as ductal cells consistent with ADM, sending signals to proliferating epithelial cells and receiving signals from myCAFs with an autocrine interaction (**Fig. 10N**). Cells in cluster 3 also showed a strong interaction with a cell neighborhood comprised of iCAFs and Schwann cells (cluster 0, **Fig. 10N**). Incoming and outgoing signaling patterns for cell clusters 0-27 were identified across all samples with diagnosed IPMN lesions (**Fig. 10O, P**). The 30 highest expressing genes across the cell clusters identified are reported in **Supplemental Figure 3**. The 30 highest expressing genes for each disease stage are reported in **Supplemental Figures 4, 5, and 6** for AC, IPMN and PDAC tissues respectively.

Collectively, these findings support the hypothesis that CD44v9+ CSCs arise in pre-malignant ADM lesions adjacent to tumors, mirroring the persistent CD44 expression observed in primary and metastatic sites by across disease stages. This suggests that CD44v9+ cells may serve as reservoirs for therapeutic resistance and recurrence. For whole tissue scans, spatial and niche maps of all biospecimens analyzed refer to the following link: https://zavros-lab.github.io/pdac-gallery/.

### Minerva Interactive Data for biospecimens 170 & 211 tumor and 170 & 211 matched adjacent cancer tissues, 009, 010 and TMA PA242f can be found using the following links

**009**: https://images.rarecyte.com/Q185-7-PDAC-009v1/#s=0#w=0#g=0#m=-1#a=-100_-100#v=4.0106_1.065_0.4143#o=-100_-100_1_1#p=Q#r=218

**170**: https://images.rarecyte.com/Q185-8-R230170/#s=1#w=6#g=7#m=-1#a=-100_-100#v=0.4747_0.5027_0.4926#o=-100_-100_1_1#p=Q#r=40

**PA242f:** https://uga-som-imtm.github.io/PA242f/

**009_010:** https://uga-som-imtm.github.io/009_010/

**170-8T-211-09T:** https://uga-som-imtm.github.io/170-8T-211-09T/

**170-09N-211-11N:** https://uga-som-imtm.github.io/170-09N-211-11N/

### CAFs persist in PDO co-cultures and recapitulate TME heterogeneity

Our research team has established an approach by which PDOs generated from surgical biospecimens collected from patients diagnosed with PDAC are successfully co-cultured with autologous immune cells (PDO/IMM co-cultures)^14^. Using this *in vitro* PDO/IMM co-culture model we demonstrated that cabozantinib depleted PMN-MDSC, and when used in combination with immune checkpoint blockade induced tumor cell death^14^. We optimized the protocol for the generation of robust co-cultures from endoscopic ultrasound guided (EUS) fine needle aspirate (FNA) (**Fig. 11A**), surgical biospecimens (**Fig. 11B**) or core biopsies of liver and lung metastatic lesions (**Fig. 11C**). MxIF revealed that PDOs generated from an FNA, prior to SOC, comprised of cell populations that heterogeneously expressed CD44v9 positive and negative tumor cells, and SPP1+ cells (**Fig. 11D, E**). In contrast to PDOs generated from FNAs, cultures derived from patients with refractory, liver metastatic PDAC were enriched for CD44v9 expressing tumor cells and SMA+ CAFs (**Fig. 11F, G**). At the time of tissue collection, blood was obtained from informed consenting patients and PBMCs generated and used for the culture of dendritic cells (DCs), cytotoxic T lymphocytes (CTLs) and myeloid derived suppressor cells (MDSCs) and used in co-cultures with autologous PDOs (**Fig. 11H**).

**Figure 11:**
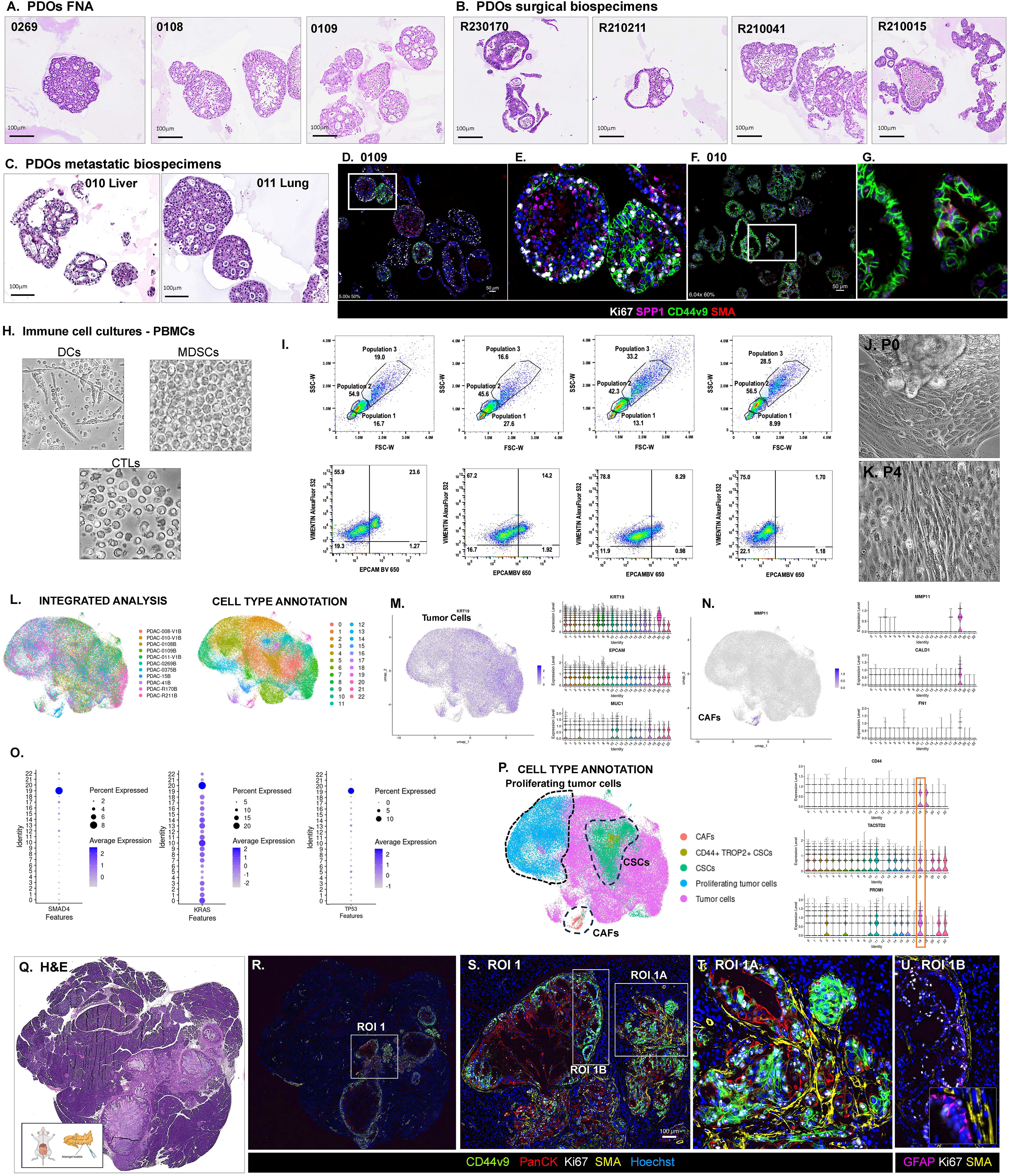
Characterization of PDO/CAF/IMM cell co-cultures. H&E staining of FFPE sections of actively growing PDO cultures generated from **(A)** FNA biospecimens, or **(B)** tumor tissue collected during the Whipple procedure or from **(C)** metastatic sites. MxIF images acquired using the Orion performed on FFPE sections of embedded PDOs generated from **(D, E)** FNA (patient 109) and **(F, G)** liver metastatic site (patient 010). **(H)** Representative bright field images of cultured dendritic cells (DCs), myeloid derived suppressor cells (MDSCs), and cytotoxic T lymphocytes (CTLs) cultures from patient PBMCs. **(I)** Flow cytometric analysis of passaged PDO cultures for the detection of the enrichment of cancer associated fibroblasts (CAFs) and fibroblasts. **(J, K)** Representative brightfield images of adherent fibroblasts and CAFs and cancer cells within PDO cultures. Single cell RNA-sequencing (scRNA-seq) analysis of PDO cultures reveals gene signatures for several cell populations including CAFs. **(L)** Integrated UMAP represents analyses performed on 11 individual patient PDO lines and identified cells populations. **(M)** Feature plot highlighting the expression of genes consistent with the tumor cell populations. **(N)** Feature plot highlighting the expression of genes consistent with the CAF populations. **(O)** Expression of SMAD4, KRAS and TP53 across all identified cell cluster populations. **(P)** Integrated UMAP identifying CAFs, CD44+TROP2+ CSCs, CSC negative for CD44 and TROP2, and proliferating tumor cells, **(Q)** Representative slide scan of H&E-stained tumor tissue collected from a NOD scid gamma mouse orthotopically transplanted with PDOs generated from patient 010. **(R-U)** Immunofluorescence staining of tissue shown in **Q** using antibodies specific for the detection of CD44v9 (green), Pan-cytokeratin (PanCK, red), Ki67 (white), SMA (yellow) and GFAP (magenta). **(R)** Selected ROI1 is shown in higher magnification in **S.** Higher magnification of ROI1A and ROI1B selected in **S,** are shown in **T and U**. Scale bars shown in (D) 50μm and (S) 100 μm.

We documented an adherent cell population consistent with the identification of CAFs (**Fig. 11I-K**). Using flow cytometry analysis, a cell population within the PDO cultures was positive for the general fibroblast marker vimentin (**Fig. 11I**). When compared to the tumor cell population marked by EPCAM with each passage of the cultures, the fibroblast population was enriched, while the EPCAM positive tumor cells diminished in culture (**Fig. 11I**). Single cell RNA sequencing of PDO cultures from eleven of the matched lines confirmed the presence of tumor cells and CAFs based on known published genes specifically expressed by these cell populations (**Fig. 11M-O**). To determine that the CAFs within the PDOs were not fibroblast-like cells derived from the tumor, classical PDAC driver genes within the cell clusters were identified (**Fig. 11P**). Cluster 19 was the main site of SMAD4 and TP53 expression, where- as KRAS expression was lowest supporting that these cells were indeed CAFs and not fibroblast-like tumor cells such as those undergoing epithelial-to-mesenchymal transition (EMT) (**Fig. 11P**). In contrast to cluster 19, tumor cell populations predominantly expressed KRAS, and cancer stem cell markers CD44 and TROP2 (**Fig. 11P**). Among the identified CAF population, cell types including cancer stem cells marked by CD44v9, TACSTD2 (TROP2) and PROM1 (CD133) were identified predominantly in cluster 18 (**Fig. 11Q**).

Orthotopically transplanted PDOs that were generated from a patient with metastasis to the liver (patient 010) were analyzed using the human-specific 14-plex Orion panel (**Fig. 11R-V**). The engrafted cells differentiated into human cancer cells expression CD44v9 (green) and CAF-associated marker SMA (yellow) (**ROI 1A, Fig. 11S-U**). Within ROI 1B we also noted the expression of GFAP supporting the engraftment of PDO Schwann cells (**Fig. 11V**). Therefore, our data show that CAFs were found to re-side within the PDO cultures that were likely carried forward from the tumor tissues at time of dissociation and persisted with processing and maintenance of cultures. These data reveal a previously unidentified cellular complexity within our group’s *in vitro* PDO/IMM co-cultures.

### Cabozantinib sensitizes PDO/CAF/IMM cell co-cultures to Pembrolizumab-induced tumor cell death, but therapy-resistant CD44v9+ cancer stem cells CD44v9 are spared

The therapeutic benefit of combinatorial therapy using cabozantinib (cabo) and Pembrolizumab (Pembro) was then studied in the preclinical PDO/CAF/IMM cell co-culture system using PDO lines generated from surgical biospecimens. Our preclinical data^14^, in combination with evidence from the prostate cancer research^15–17^ have supported the potential therapeutic benefit of pembrolizumab in combination with cabo for the treatment of PDAC. PDAC is considered a ‘non-immunogenic’ cancer via multiple mechanisms of immune evasion that include recruitment of regulatory immune cells such as T-regulatory cells (Tregs), tumor associated macrophages (TAMs) and MDCs that are known to block the anti-tumor immunity of CD8+ T cells^14,36–38^. MxIF analysis of a patient core biopsy 010 collected from a liver metastatic lesion showed expression of CD44v9+ CSCs infiltrating the surrounding liver (**Fig. 12A-C**). Analysis of ROI2 highlighted the infiltration of immunosuppressive cells expressing CD11b (MDSCs) and TAMs (CD163) (**Fig. 12D**). ROI3 showed CD8 CTLs localized predominantly within desmoplastic SMA positive stroma (**Fig. 12E**). ROI4 highlighted a histological area showing a junction of infiltrating CD44v9+ tumor cells (arrow to the left) and CD8+ CTLs localized among the hepatocytes and CD163+ Kupffer cells (arrow to the right) (**Fig. 12F, G**). Segmented CD8+ CTLs were analyzed to generate a density map that showed a ‘hotspot’ of CTL clusters (**Fig. 12H, I**) within ROI4a (area of tumor cell infiltration) and ROI4b (liver) (**Fig. 12J**). Granzyme B (GZMB)+/CD8+ cells localized pre-dominantly to the site distal to the tumor cells (ROI3 and ROI4b) (**Fig. 12J**).

**Figure 12:**
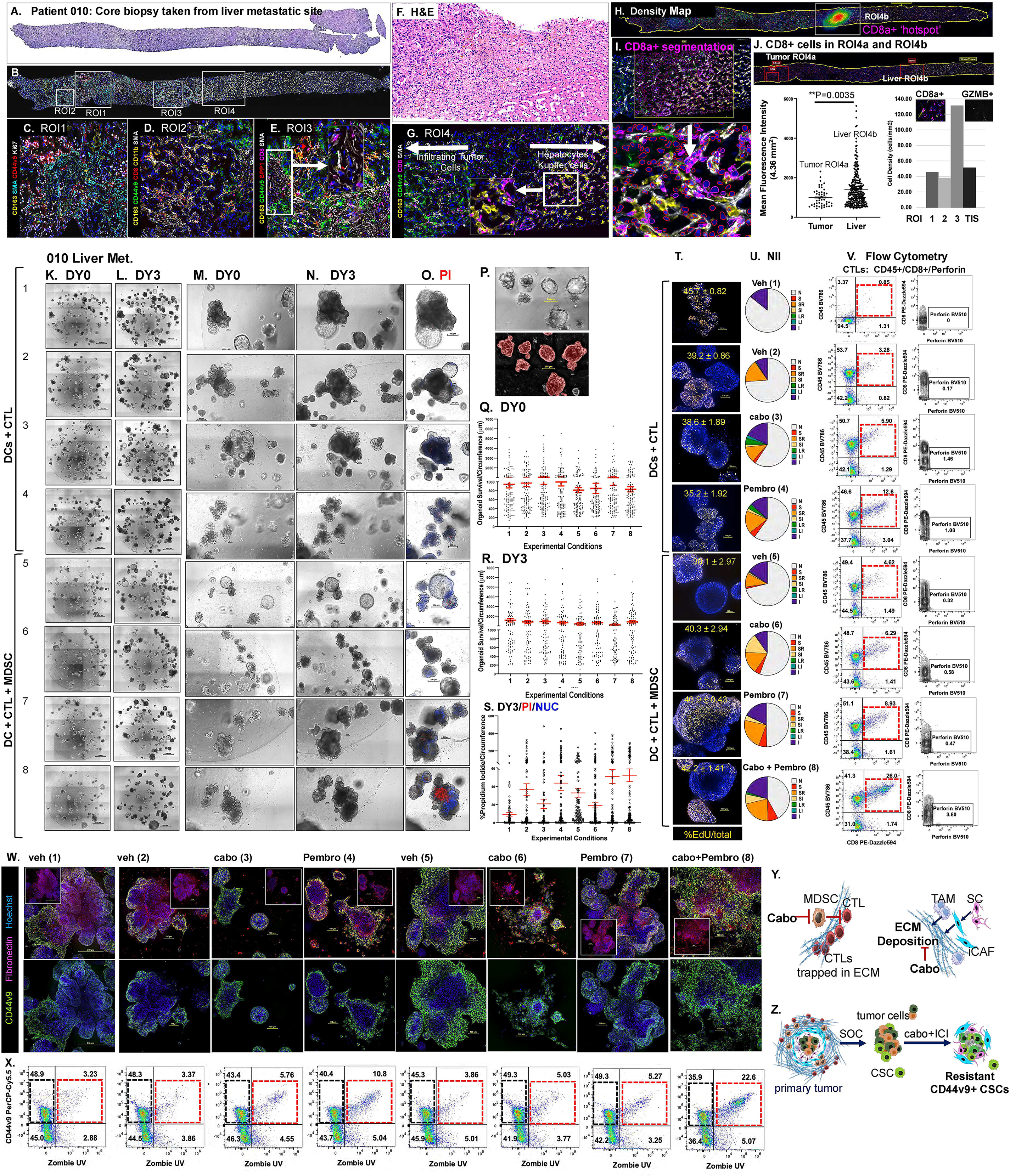
Measurement of tumor cell proliferation and viability in response to cabo plus Pembro treatment using PDO/CAF/IMM cell co-cultures generated from core biopsy of liver metastatic tumor 010. Expression of immunosuppressive cells and CD44v9 positive cancer stem cells within the metastatic PDAC TME. **(A)** Scanned H&E-stained slide of core biopsy. **(B)** Selected ROI1-4 on MxIF image. **(C)** Higher magnification of ROI1, **(D)** ROI2 and **(E)** ROI3. **(F)** ROI4 H&E stained area. **(G)** CD44v9+ (green) expressing cells and CD8+ (magenta) CTLs present among CD163 (yellow) Kupffer cells at liver metastatic site. **(H)** Density map showing CD8+ ‘hotspots’ within ROI4b. **(I)** CD8+ cell segmentation strategy used for quantification shown in J. **(J)** Measurement of the Mean Fluorescence Intensity within 4.36 mm^2^ ROI4a and ROI4b and Cell Density (cells/mm^2^) of CD8+/GZMB+ cells. **Measurement of PDO circumference. (K-N)** Images captured using phase contrast with a 10X magnification at day 0 (DY0) and day 3 (DY3) post-treatment within each well of the PD/CAF/IMM cell co-cultures. **(O)** Higher magnification and propidium iodide (PI, DY3) uptake is shown in panel for conditions 1-8. **(P)** Representative image of PDO segmentation used for calculation of circumference. Calculated PDO circumference of measurements made at **(Q)** DY0 and **(R)** DY3**. (S)** Calculated PDO PI uptake as a measurement of %PI/Circumference at DY3. **(T)** Immunofluorescence staining of proliferating EdU (orange) cells within PDO/CAF/DC/CTL cell co-cultures. **(U)** Calculation of nuclear irregularity index (NII) based on the morphometric analysis of the nuclei in experimental conditions 1-8. **(V)** Flow cytometric scatter plots presented as density dot or contour plots using a gating strategy for the expression of CD8+ CTLs expressing perforin. **(W)** Immunofluorescence staining of the expression of CD44v9 (green) and fibronectin (magenta) within PDO/CAF/DC/CTL cell co-cultures. **(X)** Flow cytometric scatter plots using a gating strategy for the expression of EPCAM+/PD-L1+ and EPCAM+/CD44v9+/Zombie+ cell populations. Schematic diagrams representing an overview of the multitargeted reprogramming of the PDAC TME in response to cabozantinib whereby **(Y)** cabo increases the efficacy of Pembro-induced tumor cell death by depleting MDSCs, inducing CTL proliferation and infiltration to the tumor site by remodeling the ECM through the modification of CAF function. **(Z)** Resistant CD44v9+ CSCs persist post SOC and in response to combinatorial treatment. **Experimental Conditions:** vehicle (1), PDO+DC+CTL vehicle (2), Cabo (3), Pembro (4), and compared to PDO/CAF/DC/CTL with MDSCs cell co-cultures treated with vehicle (5), Cabo (6), Pembro (7) or Cabo plus Pembro (8).

PDO/CAF/IMM cell co-cultures were generated from the patient’s matched tissue and used in experimental conditions designed to test the effect of cabo on MDSCs viability and CTL proliferation and effector function **(Fig. 12K-S**). Brightfield images of the same entire culture wells were acquired over a 3-day experiment (**Fig. 12K-N**) and used to measure PDO circumference (**Fig. 12P**). Over the 3-day experiment continued growth of tumor cells within patient 010 co-cultures across all conditions was observed (**Fig. 12Q, R**). Depletion of MDSCs within the PDO/CAF/IMM cell co-cultures treated with cabo (condition 6) did not increase efficacy of Pembro-induced PDO cell death (condition 8, **Fig. 12Q, R**). Phase contrast images of the same wells were acquired at day 3 of the experiment after and PI uptake was measured (**Fig. 12O, S**). Compared to vehicle controls (condition 5), PI uptake significantly increased in response to cabo plus Pembro (condition 8, **Fig. 12S**), while PDO circumference did not significantly change between DY0 (**Fig. 12Q**) and DY3 (**Fig. 12R**) in the same condition 8. Compared to vehicle controls (condition 5), PI uptake significantly increased in response to cabo plus Pembro (condition 8, **Fig. 12S**), while PDO circumference did not significantly change between DY0 (**Fig. 12Q**) and DY3 (**Fig. 12R**) in the same condition 8. While PDO proliferation was not significantly changed among conditions 2-8 (**Fig. 12T**), the NII showed an overall expansion of cells with nuclear morphology consistent with mitotic catastrophe and apoptosis, and senescence in co-culture (**Fig. 12U**). Flow cytometric analyses showed that Pembro-induced PDO apoptosis correlated with an increase in CTL proliferation and CD8+ perforin expression (**Fig. 12V**).

Immunofluorescence staining showed persistence of CD44v9+ expressing cells in response to cabo plus Pembro treatment (condition 8, **Fig. 12W**). Consistent with the immunofluorescence data, flow cytometry showed a population of CD44v9+ cells that were resistant to combinatorial cabo plus Pembro-induced tumor cell death (**Fig. 12Y**). While MDSCs blocked Pembro-induced PDO death (condition 7 compared to condition 4), cabo sensitized this response (condition 8) (**Fig. 12Y**). We also noted that cabo decreased the expression of fibronectin in co-culture conditions 3, 6 and 8 (insets shown in **Fig. 12W**). Collectively, these data show that while PDO line 010 responds to cabo plus Pembro treatment, a CSC population expressing CD44v9 was spared in co-cultures.

## DISCUSSION

In this study, we used high-plex, single cell spatial profiling and quantitatively assessed heterogeneity within limited PDAC tumor tissues and matched *in vitro* PDO/CAF/IMM cell co-cultures to demonstrate that 1) cabozantinib (cabo) increases the efficacy of Pembro-induced tumor cell death by depleting MDSCs, inducing CTL proliferation and effector function and PDO mitotic catastrophe, and 2) cabo may induce CTL infiltration to the tumor site by remodeling the ECM through the modification of CAF function. Using *in vitro* PDO/CAF/IMM cell co-cultures and matched patient tissue samples, we have also identified a tumor cell population expressing CD44v9 that is resistant to SOC chemotherapy and may attribute to recurrence of disease in patients with PDAC.

Consistent with previous reports by our group and others ^17,39–41^, we report that cabozantinib depleted the MDSCs within the TME as a mechanism to increase the efficacy of Pembrolizumab. Tumors express immune checkpoint molecules such as programmed cell death 1 ligand (PD-L1) as a mechanism to inhibit CD8+ cytotoxic T lymphocyte proliferation, survival and effector function ^42–44^. Although anti-PD1 antibodies have already been tested in clinical trials for pancreatic cancer treatment, patients have failed to respond. Of relevance to this study, MDSCs are known to block CD8+ T cell anti-tumor activity through L-arginine and L-cysteine sequestration as well as production of reactive oxygen species (ROS) ^45^ ^46^ ^47^. In a study of circulating and splenic immune cells, PDAC patients that exhibited reduced levels of CD8+ T cells also exhibited elevated levels of MDSCs ^48^. In support of the immunosuppressive MDSC function in pancreatic cancer, preclinical mouse models have similarly demonstrated that targeting the MDSC population enables an endogenous T cell response in PDAC ^49^. In humans and mice two distinct subset of MDSCs exist, monocytic-MDSCs (M-MDSCs) and polymorphonucler-MDSCs (PMN-MDSCs) ^46^ ^47^. PMN-MDSCs are known to be associated with a poor prognosis in pancreatic cancer ^49^, yet pre-clinical studies that evaluate the efficacy of targeting MDSCs in combination with chemotherapy and immunotherapy have not been performed in PDAC. Evidence from the prostate, renal and breast cancer research fields, and clinical trials, has clearly documented that tyrosine kinase inhibitors (such as cabozantinib and sunitinib) target MDSC function or generation ^15–17^. In fact, cabozantinib-targeted MDSC depletion used in combination with immune check-point blockade induces anti-tumor activities and tumor regression in castration-resistant prostate cancer ^17^. Our preclinical studies support the design for an ongoing Phase II Trial evaluating the safety and efficacy of atezolizumab in combination with cabozantinib for the treatment of metastatic, refractory PDAC (NCT04820179).

Cabozantinib consistently induced significant mitotic catastrophe that was accompanied by reduced organoid circumference and proliferation within PDO/CAF/IMM cell co-cultures. Mitotic catastrophe is known to drive either apoptosis, necrosis, or autophagy, an event preceding cell death whereby the outcome depends on the molecular profile of the cell ^50–53^. Mitotic catastrophe, which occurs under dysregulated mitosis, represents a possible strategy to specifically target tumor cells ^50–53^. While there are several anticancer drugs that are known to induce mitotic catastrophe by targeting microtubules, spindle assembly checkpoint kinases, and the DNA damage response ^50–53^, the effect of Cabozantinib has only been reported in the context of head and neck cell squamous cell carcinoma (HNSCC) ^54^. Using HNSCC cell lines, a zebra fish metastatic tumor model and mouse xenografts, Cabozantinib was found to decrease cancer cell migration, invasion and proliferation and induce mitotic catastrophe leading to apoptotic call death in both naïve, radiotherapy- and cisplatin-resistant HNSCC cells^54^. Despite the benefits of immune checkpoint inhibitors, their use as a monotherapy in PDAC, and other advanced solid tumors including HNSCC, has been disappointing and inconclusive. In support of our findings, Cabozantinib in combination with checkpoint inhibitors including pembrolizumab have proven to increase the efficacy of immunotherapy in several cancers including prostate cancer ^55,56^, renal cell carcinoma ^57^, and urothelial carcinoma ^57–61^.

Spatial transcriptomic and MxIF data revealed a population of CSCs persisting post-SOC treatment that expressed variant isoform of Cluster of Differentiation (CD) 44 transmembrane glycoprotein (CD44v), CD44v9. To our knowledge, our studies are the first to report persistence of CD44v9 with progression of disease. Studies of solid tumors show that disease recurrences frequently arise from a population of residual cancer cells often referred to as minimal residual disease^62,63^. Tumor-cell-intrinsic pathways and the TME of residual tumors regulate residual cancer cell survival and recurrence^63^. Oncogenic-induced metabolic changes lead to increased generation of mitochondrial reactive oxygen species (ROS). Subsequently there is an upregulation of antioxidant systems to balance ROS. Among these antioxidant systems is the mechanism regulated by cell adhesion molecule CD44, that suppresses the production of ROS, resulting in the therapeutic resistance and recurrence of tumors^64–69^. Alternative mRNA splicing of CD44 generates variant isoforms (CD44v) CD44v9 and CD44v6^70–72^. Among the ten CD44 variants, CD44v9 is expressed in several cancers including gastric^73^, hepatocellular^74^, pancreatic^75^, bladder^76^ and esophageal^77^. Given that CD44v9 is a known regulator of tumor cell proliferation, metastasis, and therapy resistance^78^ our findings are of relevance to the identification of the cellular origin of disease recurrence and therapy-resistance. Noteworthy was the expression of CD44v9 within acinar-to-ductal metaplastic (ADM) regions. ADM is a precursor lesion in PDAC that contributes to early neoplastic transformation, and relies on the downregulation of acinar-specific transcription factors Ptf1a, Mist1 and Nr5a2, as well as the expression of digestive enzymes such as carboxypeptidase and amylase, leading to co-expression of ductal markers (SOX2, Hnf1b, Hnf6, Pdx1, CA19-9, CAII, CD133 and osteopontin)^79,80^. Our findings support the hypothesis that CD44v9+ CSCs arise in pre-malignant ADM lesions adjacent to tumors, mirroring the persistent CD44 expression observed in primary and metastatic sites. This suggests that CD44v9+ cells may serve as reservoirs for therapeutic resistance and recurrence.

Spatial mapping for the expression of mucins MUC1, 5AC and 6 revealed that while expression of MUC5AC was not significantly different among the 3 stages of disease (AC, IPMN and PDAC), MUC1 and MUC6 expression was significant in tissues adjacent to the tumor. The expression pattern of these mucins overlapped with the expression of CD44 transcripts and specifically cancer stem cell marker CD44v9. Mucin expression patterns are often followed as a criterion for classification of precancerous lesions including PanIN and IPMN ^34^. In addition, it is reported that loss of transcription factor MIST 1 leads to enhanced PanIN development, whereby ADM is a highly plastic process involving transdifferentiation and reduced expression of acinar-specific genes including MIST1 ^35^. Indeed, MIST1 expression was significantly higher in AC tissues compared to IPMN lesions and PDAC. Network motif analysis showed that ductal-like cells consistent with ADM were found within a cellular neighborhood that exhibited an interaction with surrounding myCAFs, iCAFs and Schwann cells. Therefore, our findings support the hypothesis that CD44v9+ CSCs may arise in pre-malignant during ADM adjacent to tumors and persisting through PanIN and IPMN precancer lesions and serving as reservoirs for disease recurrence.

Data reported by PDO scRNA-seq, orthotopic transplantation and secretion of fibronectin support the existence of resident CAFs within these cultures. The decrease in fibronectin in response to cabozantinib alone and in combination with Pembrolizumab is of significance. ECM is one of the major components of PDAC that regulates mechanical support of the TME and is a source of signaling molecules that alter the characteristics of cancer cells ^81^. ECM proteins derived from tumor cells are known to reduce the efficacy of therapy and contribute to disease progression and metastasis of PDAC ^82^. Found abundantly within the PDAC ECM, cancer-associated fibroblasts (CAFs) are targets for novel anti-tumor therapy ^82–84^. Within the PDAC stroma CAFs secrete ECM-related proteins including collagen, hyaluronan (HA), and fibronectin ^85,86^, and various paracrine factors that promote tumor invasion, metastasis, TME remodeling and therapy resistance ^87^. CAFs also regulate the tumor immune microenvironment (TIME) by inducing infiltration and differentiation of regulatory T cells (T-regs), MDSCs), and tumor-associated macrophages (TAMs) ^49,88,89^, and inhibiting the cytotoxic activity of CD8^+^ cells (CTLs) in PDAC, that subsequently results in poor immunotherapy outcomes ^89–91^. CAFs are made up of heterogeneous subtypes that either promote or inhibit tumor growth ^92,93^ and the shift in phenotype is controlled by the local TME ^81–84^. Therefore, we developed a PDO culture with an identified cellular complexity that includes resident CAFs and the opportunity to add the patient’s own immune cells in co-culture. We have optimized and advanced this approach to identify *in vitro* tumor behavior with matched pathology and spatial transcriptomic signatures using small and limited PDAC patient biospecimens and blood.

Among the cell types identified within the tumor tissue transcriptomic niche and MxIF analyses were Schwann cells. Our studies presented here and previously published reports ^94–97^, using spatial biology technology, have enabled the identification for Schwann cells within the TME. The roles of cells from nerves, such as the tumor-associated/activated non-myelinating Schwann cells (TASCs) within the PDAC TME are often not considered and largely understudied. However there are key reports within the literature that document the several functions of TASCs within the TME that include regulation of the neuro-stroma niche^96^, promotion of cancer cell migration and metastasis^94^, and regulation of the tumor immune microenvironment^95,98,99^. For example, in an elegantly executed experiment using an *in vitro* 3D model, Deborde *et. al.*^94^ revealed that cancer cell invasion is mediated by a coordinated collection of TASCs that self-organize into tracks. In another study using single-cell RNA-sequencing and microarray-based spatial transcriptomic analyses, suggested that Schwann cells drive tumorigenesis by shifting the CAF function to a pro-malignant and inflammatory phenotype (iCAFs)^96^. Another study using the tamoxifen-inducible KPC mouse model suggested that TASCs within the PDAC promote an immune-resistant metabolic TME. In the same study, the depletion of TASCs expressing long noncoding RNA (lncRNA) PVT1, resulted in the suppression of tumor growth and increased efficacy of immunotherapy^97^. Collectively, these studies suggest that strategies that target the Schwann cells show promise for increasing the immunotherapy in patients with PDAC^97^.

The availability of high-plex, single cell spatial profiling and optimization of our team’s established organoid technology has allowed us to advance our prior research^14^. Taking an integrative spatial and organoid-based study approach has demonstrated that cabozantinib can remodel the PDAC TME and potentiate PD-1 immunotherapy in pre-clinical models, while resistance associates with a CD44v9+ CSC population, revealing a potential therapeutic target.

## Supporting information

file:///Users/yanazavros/Desktop/Supplemental%20Figures_Material.html

## ACKNOWLEDGEMENTS

Funding: The research team strives to translate scientific discovery to patients. We wholeheartedly thank the patients who consented to donate tumor tissues and blood for the development of the organoid cultures and spatial biology analyses. Without their willingness to participate in the study, this work would not be possible. We would like to acknowledge the support from the Merck MISP Program (Zavros), Goldman Pancreatic Cancer Grant (Zavros, University of Cincinnati), American Cancer Society Mission Boost Stage I Award (Zavros) and the National Health and Medical Research Council of Australia (Grant #2019853, Hollande, Zavros). Research reported in this study was partly supported by funds made available to Zavros through the Georgia Research Alliance and the University of Georgia School of Medicine. We acknowledge the Reaumond Foundation (Shroff) for partly supporting the CosMx analyses. The research studies were also supported by the University of Arizona Shared Resources Tissue Acquisition and Cellular/Molecular Analysis Shared Resource (TACMASR), Tissue Acquisition and Repository for Gastrointestinal and Hepatic Systems (TARGHETS, director Dr. Juanita Merchant), the Flow Cytometry and Human Immune Monitory Shared Resource (FCIMSR, director Dr. Sara Centuori), Spatial Biology and Integrative Genomics Shared Resource (SBIG, UGA School of Medicine Research Institute) and the National Cancer Institute of the National Institutes of Health under award number P30 CA023074 (University of Arizona Cancer Center).

## Author Contributions

**Pritha Adhikary:** Writing - original draft, Conceptualization, Investigation, Writing - review & editing, Methodology, Data curation, Validation, Formal analysis, Visualization

**Jayati Chakrabarti:** Writing - original draft, Conceptualization, Investigation, Writing - review & editing, Methodology, Validation, Formal analysis, Software, Visualization

**Saptarshi Mallick:** Writing - review & editing, Methodology, Resources, Data curation, Validation, Formal analysis, Software

**Xi Sun:** Investigation, Writing - review & editing, Methodology, Resources, Data curation, Validation, Formal analysis, Software, Visualization

**Ugonna Ezuma-Igwe:** Writing - original draft, Conceptualization, Investigation, Writing - review & editing, Formal analysis, Visualization

**Abigail G. Branch:** Investigation, Writing - review & editing, Resources

**Jiang Wang:** Conceptualization, Writing - review & editing, Formal analysis, Visualization

**Nicolas Braissand:** Investigation, Resources, Formal analysis, Visualization

**Megan P. Corbett:** Writing - review & editing, Formal analysis, Visualization

**Sangmin Kim:** Writing - original draft, Investigation, Writing - review & editing

**Rohan Kanakamedala:** Writing - original draft, Investigation, Writing - review & editing

**Willian C. Bastian:** Writing - review & editing, Methodology, Data curation, Validation, Formal analysis, Software, Visualization

**Eugene Douglass:** Writing - review & editing, Methodology, Data curation, Validation, Formal analysis, Software, Visualization

**Meenu Priya Resmi:** Methodology, Formal analysis

**Payton Stevens:** Conceptualization, Writing - review & editing, Methodology

**Marco Padilla-Rodriguez:** Conceptualization, Writing - review & editing, Resources

**Maga Sanchez:** Methodology, Validation

**Gregory L. Beatty:** Conceptualization, Writing - review & editing

**Elliott B. Burdette:** Conceptualization, Writing - review & editing, Formal analysis, Visualization

**James Griffin:** Conceptualization, Writing - review & editing, Visualization

**Petros G. Nikolinankos:** Conceptualization, Writing - review & editing, Visualization

**Kellen Chen:** Writing - review & editing, Resources, Data Curation

**Taylor Riall:** Conceptualization, Writing - review & editing, Resources, Data Curation

**Syed A. Ahmad:** Conceptualization, Writing - review & editing, Resources

**Rachna Shroff:** Conceptualization, Writing - review & editing, Resources

**Frédéric Hollande:** Conceptualization, Writing - review & editing, Methodology, Funding acquisition, Supervision

**Davendra Sohal:** Writing - original draft, Investigation, Writing -review & editing, Methodology, Data curation, Validation

**Yana Zavros:** Writing - original draft, Conceptualization, Investigation, Writing - review & editing, Methodology, Resources, Funding acquisition, Data curation, Validation, Supervision, Formal analysis, Project administration, Visualization We would also like to thank Melinda Duplessis, Edward Lo, Robert Wilson, Tad George, Jon Ladd, Alexis Coffer and the entire the Rarecyte team for their continued technical support and guidance. Finally, we thank UGA’s School of Medicine and College of Veterinary Medicine IT teams in particular Adam Evans, Kevin Whiten and Matthew Blankenship for their assistance with data management and storage.

## Data, Code, and Materials Availability

All data and code needed to evaluate and reproduce the results in the paper are present in the paper and/or the Supplementary Materials. MTA has been provided in Supplementary Material. The data and codes needed can be provided by the research team pending scientific review and a completed material transfer agreement. Requests for the data should be submitted to

## TABLE OF CONTENTS: SUPPLEMENTARY MATERIAL

**Supplementary Table 1:** Patient Clinical Data of Collected Biospecimens.

**Supplementary Table 2:** Spectral Flow Cytometry (Cytek™ Aurora) Antibody Panel

**Supplementary Figure 1:** Cytek®Full Spectrum Viewer and Calculation of Similarity Index

**Supplementary Figure 2:** The top differentially expressed genes were identified for each cell type identified on the UMAP.

**Supplementary Figure 3:** The top differentially expressed genes were reported for each cell type identified in tissues analyzed by CosMx whole transcriptomic (WTx) SMI approach.

**Supplementary Figure 4:** The 30 highest expressing genes are reported for adjacent cancer (AC) tissues analyzed by CosMx whole transcriptomic (WTx) SMI.

**Supplementary Figure 5:** The 30 highest expressing genes are reported for IPMN tissues analyzed by CosMx whole transcriptomic (WTx) SMI.

**Supplementary Figure 6:** The 30 highest expressing genes are reported for PDAC tissues analyzed by CosMx whole transcriptomic (WTx) SMI.

**Supplemental Figure 7: Spatial proximity of CD44v9**⁺ **cells to immune and stromal cell populations in PDAC tissue. (A)** Representative MxIF image of biospecimen 010 collected from a liver metastatic lesion of a patient with PDAC showing regions of interest (ROIs) selected for spatial analysis. **(B)** Percentage of total cells positive for SMA, CD11b, CD8, and CD163 within ROI1 and ROI4. **(C–K)** HALO proximity maps showing the spatial distribution and proximity of CD44v9⁺ cells to CD11b⁺ **(C, D)**, CD8⁺ **(F, G)**, CD163⁺ **(H, I)**, and SMA⁺ **(J, K)** cells in ROI1 and ROI4. Proximity lines indicate spatial distance between CD44v9⁺ cells and the indicated cell population. **(L–S)** Representative multiplex immunofluorescence images corresponding to the proximity analyses, showing CD44v9 and CD11b **(L, M)**, CD8 **(N, O)**, CD163 **(P, Q)**, or SMA **(R, S)** expression in ROI1 and ROI4. **(T–W)** Quantification of CD8⁺ **(T)**, CD163⁺ **(U)**, CD11b⁺ **(V)**, and SMA⁺ **(W)** cells.

## REFERENCES

1 Ercan, G., Karlitepe, A. & Ozpolat, B. Pancreatic Cancer Stem Cells and Therapeutic Approaches. Anticancer Res 37, 2761–2775 (2017). 10.21873/anticanres.11628

2 Fitzgerald, T. L. & McCubrey, J. A. Pancreatic cancer stem cells: association with cell surface markers, prognosis, resistance, metastasis and treatment. Adv Biol Regul 56, 45–50 (2014). 10.1016/j.jbior.2014.05.001

3 Quante, M., Varga, J., Wang, T. C. & Greten, F. R. The gastrointestinal tumor microenvironment. Gastroenterology 145, 63–78 (2013). 10.1053/j.gastro.2013.03.052

4 Hosein, A. N., Brekken, R. A. & Maitra, A. Pancreatic cancer stroma: an update on therapeutic targeting strategies. Nat Rev Gastroenterol Hepatol (2020). 10.1038/s41575-020-0300-1

5 Atkins, M. B. & Tannir, N. M. Current and emerging therapies for first-line treatment of metastatic clear cell renal cell carcinoma. Cancer Treat Rev 70, 127–137 (2018). 10.1016/j.ctrv.2018.07.009

6 Choueiri, T. K. et al. Cabozantinib versus Everolimus in Advanced Renal-Cell Carcinoma. N Engl J Med 373, 1814–1823 (2015). 10.1056/NEJMoa1510016

7 Abou-Alfa, G. K. et al. Cabozantinib in Patients with Advanced and Progressing Hepatocellular Carcinoma. N Engl J Med 379, 54–63 (2018). 10.1056/NEJMoa1717002

8 Schlumberger, M. et al. Overall survival analysis of EXAM, a phase III trial of cabozantinib in patients with radiographically progressive medullary thyroid carcinoma. Ann Oncol 28, 2813–2819 (2017). 10.1093/annonc/mdx479

9 Fukumura, D., Kloepper, J., Amoozgar, Z., Duda, D. G. & Jain, R. K. Enhancing cancer immunotherapy using antiangiogenics: opportunities and challenges. Nat Rev Clin Oncol 15, 325–340 (2018). 10.1038/nrclinonc.2018.29

10 Santoni, M. et al. Antitumor effects of the multi-target tyrosine kinase inhibitor cabozantinib: a comprehensive review of the preclinical evidence. Expert Rev Anticancer Ther 21, 1029–1054 (2021). 10.1080/14737140.2021.1919090

11 Gherardi, E., Birchmeier, W., Birchmeier, C. & Vande Woude, G. Targeting MET in cancer: rationale and progress. Nat Rev Cancer 12, 89–103 (2012). 10.1038/nrc3205

12 Graham, D. K., DeRyckere, D., Davies, K. D. & Earp, H. S. The TAM family: phosphatidylserine sensing receptor tyrosine kinases gone awry in cancer. Nat Rev Cancer 14, 769–785 (2014). 10.1038/nrc3847

13 Zhen, D. B. et al. A phase I trial of cabozantinib and gemcitabine in advanced pancreatic cancer. Invest New Drugs 34, 733–739 (2016). 10.1007/s10637-016-0376-1

14 Holokai, L. et al. Murine- and Human-Derived Autologous Organoid/Immune Cell Co-Cultures as Pre-Clinical Models of Pancreatic Ductal Adenocarcinoma. Cancers (Basel) 12 (2020). 10.3390/cancers12123816

15 Ko, J. S. et al. Sunitinib mediates reversal of myeloid-derived suppressor cell accumulation in renal cell carcinoma patients. Clin Cancer Res 15, 2148–2157 (2009). 10.1158/1078-0432.CCR-08-1332

16 Kodera, Y. et al. Sunitinib inhibits lymphatic endothelial cell functions and lymph node metastasis in a breast cancer model through inhibition of vascular endothelial growth factor receptor 3. Breast Cancer Res 13, R66 (2011). 10.1186/bcr2903

17 Lu, X. et al. Effective combinatorial immunotherapy for castration-resistant prostate cancer. Nature 543, 728–732 (2017). 10.1038/nature21676

18 Chakrabarti, J. et al. Mouse-Derived Gastric Organoid and Immune Cell Co-culture for the Study of the Tumor Microenvironment. Methods Mol Biol 1817, 157–168 (2018). 10.1007/978-1-4939-8600-2_16

19 Filippi-Chiela, E. C. et al. Nuclear morphometric analysis (NMA): screening of senescence, apoptosis and nuclear irregularities. PLoS One 7, e42522 (2012). 10.1371/journal.pone.0042522

20 Steele, N. G. et al. An Organoid-Based Preclinical Model of Human Gastric Cancer. Cell Mol Gastroenterol Hepatol 7, 161–184 (2019). 10.1016/j.jcmgh.2018.09.008

21 Hao, Y. et al. Dictionary learning for integrative, multimodal and scalable single-cell analysis. Nat Biotechnol 42, 293–304 (2024). 10.1038/s41587-023-01767-y

22 Patikas, N. et al. Integration of large, complex single-cell datasets with Harmony2. bioRxiv (2026). 10.64898/2026.03.16.711825

23 Steele, N. G. et al. Multimodal Mapping of the Tumor and Peripheral Blood Immune Landscape in Human Pancreatic Cancer. Nat Cancer 1, 1097–1112 (2020). 10.1038/s43018-020-00121-4

24 Chen, K. et al. Development and validation of prognostic and diagnostic model for pancreatic ductal adenocarcinoma based on scRNA-seq and bulk-seq datasets. Hum Mol Genet 31, 1705–1719 (2022). 10.1093/hmg/ddab343

25 Lin, W. et al. Single-cell transcriptome analysis of tumor and stromal compartments of pancreatic ductal adenocarcinoma primary tumors and metastatic lesions. Genome Med 12, 80 (2020). 10.1186/s13073-020-00776-9

26 Cui Zhou, D., et al. Spatially restricted drivers and transitional cell populations cooperate with the microenvironment in untreated and chemo-resistant pancreatic cancer. Nat Genet 54, 1390–1405 (2022). 10.1038/s41588-022-01157-1

27 Wang, Y. et al. Single-cell analysis of pancreatic ductal adenocarcinoma identifies a novel fibroblast subtype associated with poor prognosis but better immunotherapy response. Cell Discov 7, 36 (2021). 10.1038/s41421-021-00271-4

28 Ianevski, A., Giri, A. K. & Aittokallio, T. Fully-automated and ultra-fast cell-type identification using specific marker combinations from single-cell transcriptomic data. Nature Communications 13, 1246 (2022). 10.1038/s41467-022-28803-w

29 Kushima, R. The updated WHO classification of digestive system tumours-gastric adenocarcinoma and dysplasia. Pathologe 43, 8–15 (2022). 10.1007/s00292-021-01023-7

30 Nagtegaal, I. D. et al. The 2019 WHO classification of tumours of the digestive system. Histopathology 76, 182–188 (2020). 10.1111/his.13975

31 Bankhead, P. et al. QuPath: Open source software for digital pathology image analysis. Sci Rep 7, 16878 (2017). 10.1038/s41598-017-17204-5

32 Choudhary, S. & Satija, R. Comparison and evaluation of statistical error models for scRNA-seq. Genome Biol 23, 27 (2022). 10.1186/s13059-021-02584-9

33 Werba, G. et al. Single-cell RNA sequencing reveals the effects of chemotherapy on human pancreatic adenocarcinoma and its tumor microenvironment. Nat Commun 14, 797 (2023). 10.1038/s41467-023-36296-4

34 Wang, S., You, L., Dai, M. & Zhao, Y. Mucins in pancreatic cancer: A well-established but promising family for diagnosis, prognosis and therapy. J Cell Mol Med 24, 10279–10289 (2020). 10.1111/jcmm.15684

35 Schneider, A. T. et al. A decision point between transdifferentiation and programmed cell death priming controls KRAS-dependent pancreatic cancer development. Nat Commun 16, 1765 (2025). 10.1038/s41467-025-56493-7

36 Carstens, J. L. et al. Spatial computation of intratumoral T cells correlates with survival of patients with pancreatic cancer. Nat Commun 8, 15095 (2017). 10.1038/ncomms15095

37 Attri, K. S., Mehla, K. & Singh, P. K. Evaluation of Macrophage Polarization in Pancreatic Cancer Microenvironment Under Hypoxia. Methods Mol Biol 1742, 265–276 (2018). 10.1007/978-1-4939-7665-2_23

38 Wartenberg, M. et al. Accumulation of FOXP3+T-cells in the tumor microenvironment is associated with an epithelial-mesenchymal-transition-type tumor budding phenotype and is an independent prognostic factor in surgically resected pancreatic ductal adenocarcinoma. Oncotarget 6, 4190–4201 (2015). 10.18632/oncotarget.2775

39 Patnaik, A. et al. Cabozantinib Eradicates Advanced Murine Prostate Cancer by Activating Antitumor Innate Immunity. Cancer Discov 7, 750–765 (2017). 10.1158/2159-8290.CD-16-0778

40 Liu, H. et al. Tyrosine Kinase Inhibitor Cabozantinib Inhibits Murine Renal Cancer by Activating Innate and Adaptive Immunity. Front Oncol 11, 663517 (2021). 10.3389/fonc.2021.663517

41 Khaki Bakhtiarvand, V., et al. Myeloid-derived suppressor cells (MDSCs) depletion by cabozantinib improves the efficacy of anti-HER2 antibody-based immunotherapy in a 4T1-HER2 murine breast cancer model. Int Immunopharmacol 113, 109470 (2022). 10.1016/j.intimp.2022.109470

42 Ahmadzadeh, M. et al. Tumor antigen-specific CD8 T cells infiltrating the tumor express high levels of PD-1 and are functionally impaired. Blood 114, 1537–1544 (2009). 10.1182/blood-2008-12-195792

43 Chen, X. et al. PD-1 regulates extrathymic regulatory T-cell differentiation. Eur J Immunol 44, 2603–2616 (2014). 10.1002/eji.201344423

44 Reissfelder, C. et al. Tumor-specific cytotoxic T lymphocyte activity determines colorectal cancer patient prognosis. J Clin Invest 125, 739–751 (2015). 10.1172/JCI74894

45 Zhang, Y. et al. Myeloid cells are required for PD-1/PD-L1 checkpoint activation and the establishment of an immunosuppressive environment in pancreatic cancer. Gut 66, 124–136 (2017). 10.1136/gutjnl-2016-312078

46 Schouppe, E., Van Overmeire, E., Laoui, D., Keirsse, J. & Van Ginderachter, J. A. Modulation of CD8(+) T-cell activation events by monocytic and granulocytic myeloid-derived suppressor cells. Immunobiology 218, 1385–1391 (2013). 10.1016/j.imbio.2013.07.003

47 Youn, J. I., Nagaraj, S., Collazo, M. & Gabrilovich, D. I. Subsets of myeloid-derived suppressor cells in tumor-bearing mice. J Immunol 181, 5791–5802 (2008).

48 Basso, D. et al. Pancreatic tumors and immature immunosuppressive myeloid cells in blood and spleen: role of inhibitory co-stimulatory molecules PDL1 and CTLA4. An in vivo and in vitro study. PLoS One 8, e54824 (2013). 10.1371/journal.pone.0054824

49 Stromnes, I. M. et al. Targeted depletion of an MDSC subset unmasks pancreatic ductal adenocarcinoma to adaptive immunity. Gut 63, 1769–1781 (2014). 10.1136/gutjnl-2013-306271

50 Castedo, M. et al. Cell death by mitotic catastrophe: a molecular definition. Oncogene 23, 2825–2837 (2004). 10.1038/sj.onc.1207528

51 Bunz, F. et al. Requirement for p53 and p21 to sustain G2 arrest after DNA damage. Science 282, 1497–1501 (1998). 10.1126/science.282.5393.1497

52 Andreassen, P. R., Lacroix, F. B., Lohez, O. D. & Margolis, R. L. Neither p21WAF1 nor 14-3-3sigma prevents G2 progression to mitotic catastrophe in human colon carcinoma cells after DNA damage, but p21WAF1 induces stable G1 arrest in resulting tetraploid cells. Cancer Res 61, 7660–7668 (2001).

53 Roninson, I. B., Broude, E. V. & Chang, B. D. If not apoptosis, then what? Treatment-induced senescence and mitotic catastrophe in tumor cells. Drug Resist Updat 4, 303–313 (2001). 10.1054/drup.2001.0213

54 Hagege, A. et al. Targeting of c-MET and AXL by cabozantinib is a potential therapeutic strategy for patients with head and neck cell carcinoma. Cell Rep Med 3, 100659 (2022). 10.1016/j.xcrm.2022.100659

55 Smith, D. C. et al. Efficacy and Effect of Cabozantinib on Bone Metastases in Treatment-naive Castration-resistant Prostate Cancer. Clin Genitourin Cancer 18, 332–339 e332 (2020). 10.1016/j.clgc.2019.10.019

56 Agarwal, N. et al. Cabozantinib in combination with atezolizumab in patients with metastatic castration-resistant prostate cancer: results from an expansion cohort of a multicentre, open-label, phase 1b trial (COSMIC-021). Lancet Oncol 23, 899–909 (2022). 10.1016/S1470-2045(22)00278-9

57 Choueiri, T. K. et al. Cabozantinib plus Nivolumab and Ipilimumab in Renal-Cell Carcinoma. N Engl J Med 388, 1767–1778 (2023). 10.1056/NEJMoa2212851

58 Apolo, A. B. et al. Final Results From a Phase I Trial and Expansion Cohorts of Cabozantinib and Nivolumab Alone or With Ipilimumab for Advanced/Metastatic Genitourinary Tumors. J Clin Oncol 42, 3033–3046 (2024). 10.1200/JCO.23.02233

59 Apolo, A. B. et al. Phase I Study of Cabozantinib and Nivolumab Alone or With Ipilimumab for Advanced or Metastatic Urothelial Carcinoma and Other Genitourinary Tumors. J Clin Oncol 38, 3672–3684 (2020). 10.1200/JCO.20.01652

60 Apolo, A. B. et al. Cabozantinib in patients with platinum-refractory metastatic urothelial carcinoma: an open-label, single-centre, phase 2 trial. Lancet Oncol 21, 1099–1109 (2020). 10.1016/S1470-2045(20)30202-3

61 Girardi, D. M. et al. Cabozantinib plus Nivolumab Phase I Expansion Study in Patients with Metastatic Urothelial Carcinoma Refractory to Immune Checkpoint Inhibitor Therapy. Clin Cancer Res 28, 1353–1362 (2022). 10.1158/1078-0432.CCR-21-3726

62 Zhu, L. et al. Minimal residual disease (MRD) detection in solid tumors using circulating tumor DNA: a systematic review. Front Genet 14, 1172108 (2023). 10.3389/fgene.2023.1172108

63 Ceyhan, Y., Garcia, N. M. G. & Alvarez, J. V. Immune cells in residual disease and recurrence. Trends Cancer 9, 554–565 (2023). 10.1016/j.trecan.2023.04.001

64 He, Y. et al. CD44 is overexpressed and correlated with tumor progression in gallbladder cancer. Cancer Manag Res 10, 3857–3865 (2018). 10.2147/CMAR.S175681

65 Zhang, C., Li, C., He, F., Cai, Y. & Yang, H. Identification of CD44+CD24+ gastric cancer stem cells. J Cancer Res Clin Oncol 137, 1679–1686 (2011). 10.1007/s00432-011-1038-5

66 Engevik, A. C. et al. The Development of Spasmolytic Polypeptide/TFF2-Expressing Metaplasia (SPEM) During Gastric Repair Is Absent in the Aged Stomach. Cell Mol Gastroenterol Hepatol 2, 605–624 (2016). 10.1016/j.jcmgh.2016.05.004

67 Ishimoto, T. et al. CD44 variant regulates redox status in cancer cells by stabilizing the xCT subunit of system xc(-) and thereby promotes tumor growth. Cancer Cell 19, 387–400 (2011). 10.1016/j.ccr.2011.01.038

68 Masson, D. et al. Soluble CD44: quantification and molecular repartition in plasma of patients with colorectal cancer. Br J Cancer 80, 1995–2000 (1999). 10.1038/sj.bjc.6690633

69 Guo, Y. J. et al. Potential use of soluble CD44 in serum as indicator of tumor burden and metastasis in patients with gastric or colon cancer. Cancer Res 54, 422–426 (1994).

70 Medrano-Gonzalez, P. A., Rivera-Ramirez, O., Montano, L. F. & Rendon-Huerta, E. P. Proteolytic Processing of CD44 and Its Implications in Cancer. Stem Cells Int 2021, 6667735 (2021). 10.1155/2021/6667735

71 Zavros, Y. Initiation and Maintenance of Gastric Cancer: A Focus on CD44 Variant Isoforms and Cancer Stem Cells. Cell Mol Gastroenterol Hepatol 4, 55–63 (2017). 10.1016/j.jcmgh.2017.03.003

72 Zoller, M. CD44: can a cancer-initiating cell profit from an abundantly expressed molecule? Nat Rev Cancer 11, 254–267 (2011). 10.1038/nrc3023

73 Wada, T. et al. Functional role of CD44v-xCT system in the development of spasmolytic polypeptide-expressing metaplasia. Cancer Sci 104, 1323–1329 (2013). 10.1111/cas.12236

74 Wada, F. et al. High expression of CD44v9 and xCT in chemoresistant hepatocellular carcinoma: Potential targets by sulfasalazine. Cancer Sci 109, 2801–2810 (2018). 10.1111/cas.13728

75 Kiuchi, S., Ikeshita, S., Miyatake, Y. & Kasahara, M. Pancreatic cancer cells express CD44 variant 9 and multidrug resistance protein 1 during mitosis. Exp Mol Pathol 98, 41–46 (2015). 10.1016/j.yexmp.2014.12.001

76 Ogihara, K. et al. Sulfasalazine could modulate the CD44v9-xCT system and enhance cisplatin-induced cytotoxic effects in metastatic bladder cancer. Cancer Sci 110, 1431–1441 (2019). 10.1111/cas.13960

77 Taniguchi, D. et al. CD44v9 is associated with epithelial-mesenchymal transition and poor outcomes in esophageal squamous cell carcinoma. Cancer Med 7, 6258–6268 (2018). 10.1002/cam4.1874

78 Chen, C., Zhao, S., Karnad, A. & Freeman, J. W. The biology and role of CD44 in cancer progression: therapeutic implications. J Hematol Oncol 11, 64 (2018). 10.1186/s13045-018-0605-5

79 Jiang, J. et al. Transcriptional Profile of Human Pancreatic Acinar Ductal Metaplasia. Gastro Hep Adv 2, 532–543 (2023). 10.1016/j.gastha.2023.02.003

80 Nishimon, R. et al. Pancreatic ductal adenocarcinoma with acinar-to-ductal metaplasia-like cancer cells shows increased cellular proliferation. Pancreatology 23, 811–817 (2023). 10.1016/j.pan.2023.08.007

81 Huang, J. et al. Extracellular matrix and its therapeutic potential for cancer treatment. Signal Transduct Target Ther 6, 153 (2021). 10.1038/s41392-021-00544-0

82 Tiriac, H. et al. Organoid Profiling Identifies Common Responders to Chemotherapy in Pancreatic Cancer. Cancer Discov 8, 1112–1129 (2018). 10.1158/2159-8290.CD-18-0349

83 Dreyer, S. B. et al. Genomic and Molecular Analyses Identify Molecular Subtypes of Pancreatic Cancer Recurrence. Gastroenterology 162, 320–324 e324 (2022). 10.1053/j.gastro.2021.09.022

84 Bailey, P. et al. Genomic analyses identify molecular subtypes of pancreatic cancer. Nature 531, 47–52 (2016). 10.1038/nature16965

85 Ferrara, B. et al. The Extracellular Matrix in Pancreatic Cancer: Description of a Complex Network and Promising Therapeutic Options. Cancers (Basel) 13 (2021). 10.3390/cancers13174442

86 McCarroll, J. A. et al. Role of pancreatic stellate cells in chemoresistance in pancreatic cancer. Front Physiol 5, 141 (2014). 10.3389/fphys.2014.00141

87 Xu, Z. et al. Role of pancreatic stellate cells in pancreatic cancer metastasis. Am J Pathol 177, 2585–2596 (2010). 10.2353/ajpath.2010.090899

88 Zhang, Y. et al. CD4+ T lymphocyte ablation prevents pancreatic carcinogenesis in mice. Cancer Immunol Res 2, 423–435 (2014). 10.1158/2326-6066.CIR-14-0016-T

89 Zhu, Y. et al. Tissue-Resident Macrophages in Pancreatic Ductal Adenocarcinoma Originate from Embryonic Hematopoiesis and Promote Tumor Progression. Immunity 47, 597 (2017). 10.1016/j.immuni.2017.08.018

90 Zhang, Y. et al. Regulatory T-cell Depletion Alters the Tumor Microenvironment and Accelerates Pancreatic Carcinogenesis. Cancer Discov 10, 422–439 (2020). 10.1158/2159-8290.CD-19-0958

91 Qiu, J. et al. mTOR inhibitor, gemcitabine and PD-L1 antibody blockade combination therapy suppresses pancreatic cancer progression via metabolic reprogramming and immune microenvironment remodeling in Trp53(flox/+)LSL-Kras(G12D/+)Pdx-1-Cre murine models. Cancer Lett 554, 216020 (2023). 10.1016/j.canlet.2022.216020

92 Tsoumakidou, M. The advent of immune stimulating CAFs in cancer. Nat Rev Cancer 23, 258–269 (2023). 10.1038/s41568-023-00549-7

93 Geng, X. et al. Cancer-Associated Fibroblast (CAF) Heterogeneity and Targeting Therapy of CAFs in Pancreatic Cancer. Front Cell Dev Biol 9, 655152 (2021). 10.3389/fcell.2021.655152

94 Deborde, S. et al. Reprogrammed Schwann Cells Organize into Dynamic Tracks that Promote Pancreatic Cancer Invasion. Cancer Discov 12, 2454–2473 (2022). 10.1158/2159-8290.CD-21-1690

95 Cai, Z. et al. Schwann cells in pancreatic cancer: Unraveling their multifaceted roles in tumorigenesis and neural interactions. Cancer Lett 587, 216689 (2024). 10.1016/j.canlet.2024.216689

96 Xue, M. et al. Schwann cells regulate tumor cells and cancer-associated fibroblasts in the pancreatic ductal adenocarcinoma microenvironment. Nat Commun 14, 4600 (2023). 10.1038/s41467-023-40314-w

97 Sun, C. et al. Tumor-associated nonmyelinating Schwann cell-expressed PVT1 promotes pancreatic cancer kynurenine pathway and tumor immune exclusion. Sci Adv 9, eadd6995 (2023). 10.1126/sciadv.add6995

98 Martyn, G. V., Shurin, G. V., Keskinov, A. A., Bunimovich, Y. L. & Shurin, M. R. Schwann cells shape the neuro-immune environs and control cancer progression. Cancer Immunol Immunother 68, 1819–1829 (2019). 10.1007/s00262-018-02296-3

99 Shurin, G. V., Vats, K., Kruglov, O., Bunimovich, Y. L. & Shurin, M. R. Tumor-Induced T Cell Polarization by Schwann Cells. Cells 11 (2022). 10.3390/cells11223541

