## Supplementary material for "Spatial Biology and Organoid Technologies Reveal a Potential Therapy-Resistant Cancer Stem Cell Population in Pancreatic Ductal Adenocarcinoma": file:///Users/yanazavros/Desktop/Supplemental%20Figures_Material.html

### Supplemental Table 1: Patient Clinical Data of Collected Biospecimens.

| Patient tissue samples | Histologic Type | Collection Procedure | Staging | Site | Grade |
|---|---|---|---|---|---|
| **0108 PAA.C.1a** | Ductal adenocarcinoma | Fine needle aspiration | pT3pN0 | Pancreas | - |
| **0109 PAA.C.1a** | Ductal adenocarcinoma | Fine needle aspiration | - | Pancreas | - |
| **0269 PAF.C.1a** | Ductal adenocarcinoma | Fine needle aspiration | - | Pancreas | - |
| **0375 PAA.C.1a** | Ductal adenocarcinoma | Fine needle aspiration | pT3pN0 | Pancreas | - |
| **R210015** | Adenocarcinoma, intestinal type | Pancreaticoduodenectomy (Whipple resection) | pT1bN0 | Intra -ampullary | G2: Moderately differentiated |
| **R230170** | Ductal adenocarcinoma | Pancreaticoduodenectomy, (Whipple resection) Partial pancreatectomy | YpT2N0 | Pancreatic head | G1: Well-differentiated |
| **R210211** | Ductal adenocarcinoma | Pancreaticoduodenectomy, (Whipple resection) | pT3pN2 | Pancreatic head Distal Bile duct | G3: poorly differentiated |
| **R220019** | Neuroendocrine | Pancreaticoduodenectomy, (Whipple resection) | pT3pN0 | Pancreatic body & tail | G1: well differentiated |
| **004** | Ductal adenocarcinoma | Core biopsy | Metastasis | Liver | - |
| **006** | Ductal adenocarcinoma | Core biopsy | Metastasis | Lung | - |
| **009** | Ductal adenocarcinoma | Core biopsy | Metastasis | Liver | - |
| **010** | Ductal adenocarcinoma | Core biopsy | Metastasis | Liver | - |
| **011** | Ductal adenocarcinoma | Core biopsy | Metastasis | Lung | - |
| **015** | Ductal adenocarcinoma | Core biopsy | Metastasis | Liver | - |
| **TMA**** | Ductal adenocarcinoma, normal, Duodenal invasion | 10 PDAC<br>2 Metastatic lesions (duodenum)<br>2 Mucinous<br>4 Normal pancreas | | | |

****TMA – refer to information provided by manufacturer - TissueArray.com, catalogue numbers PA242e and PA242f**

**Supplemental Table 2: Spectral flow cytometry (Cytek™ Aurora) Antibody Panel**

| Antibody | Type | Fluorophores | Catalog Number (Primary/Conjugated) | Catalog Number (Secondary) |
|---|---|---|---|---|
| Dead | Conjugated | Zombie UV | Biolegend # 423107 | |
| CFSE | Conjugated | FITC | Biolegend # 423801 | |
| HLA-DR | Conjugated | BUV 395 | BD # 565972 | |
| CD163 | Conjugated | BV 421 | BD # 562643 | |
| Perforin | Conjugated | BV 510 | Biolegend # 308120 | |
| CD45 | Conjugated | BV 786 | BD # 563716 | |
| CD44v9 | Unconjugated | PerCp/Cy5.5 | CosmoBio # LKG-M001 | Biolegend # 405424 |
| Arg1 | Conjugated | PE | Biolegend # 369704 | |
| CD8 | Conjugated | PE Dazzle 594 | Biolegend # 344744 | |
| CD11b | Conjugated | PE/Cy5 | Biolegend # 301308 | |
| CD137 | Conjugated | BV 605 | Biolegend # 309822 | |
| CD15 | Conjugated | PE/Cy7 | Biolegend # 301924 | |
| CD14 | Conjugated | APC/Cy7 | Biolegend # 301820 | |
| PD-L1 | Conjugated | APC | Biolegend # 329707 | |
| CD33 | Conjugated | BV 785 | Biolegend # 303428 | |
| EpCAM | Conjugated | BV 650 | Biolegend # 324226 | |

#### Supplemental Figure 1: Cytek®Full Spectrum Viewer and Calculation of Similarity Index

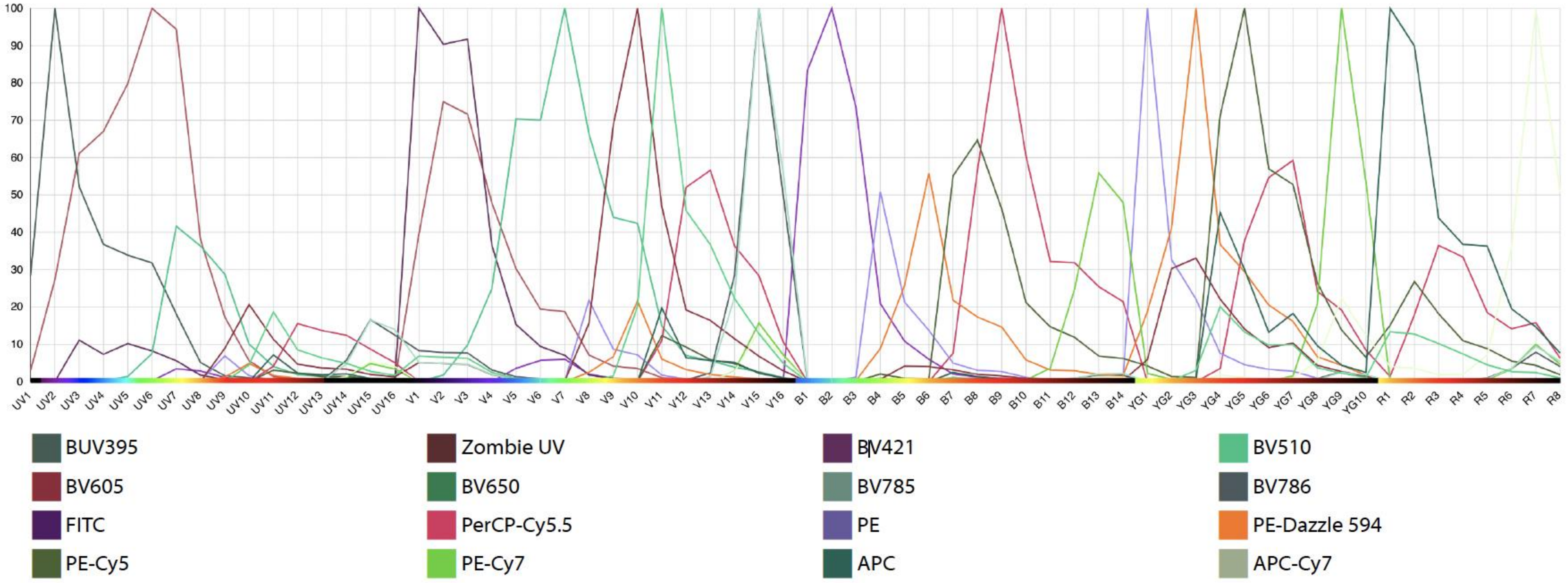


| | | | | | | | | | | | | | | | | |
|---|---|---|---|---|---|---|---|---|---|---|---|---|---|---|---|---|
| BUV395 | 1 | | | | | | | | | | | | | | | |
| Zombie UV | 0.54 | 1 | | | | | | | | | | | | | | |
| BV421 | 0.08 | 0.6 | 1 | | | | | | | | | | | | | |
| BV510 | 0.05 | 0.36 | 0.16 | 1 | | | | | | | | | | | | |
| BV605 | 0 | 0.06 | 0.06 | 0.37 | 1 | | | | | | | | | | | |
| BV650 | 0 | 0.06 | 0.09 | 0.15 | 0.53 | 1 | | | | | | | | | | |
| BV786 | 0 | 0.08 | 0.12 | 0.04 | 0.08 | 0.17 | 1 | | | | | | | | | |
| BV785 | 0 | 0.04 | 0.08 | 0.03 | 0.08 | 0.15 | 0.89 | 1 | | | | | | | | |
| FITC | 0.01 | 0.03 | 0.01 | 0.06 | 0.01 | 0 | 0 | 0 | 1 | | | | | | | |
| PerCP-Cy5.5 | 0 | 0 | 0 | 0.03 | 0.19 | 0.39 | 0.22 | 0.21 | 0.01 | 1 | | | | | | |
| PE | 0 | 0.01 | 0 | 0.11 | 0.25 | 0.05 | 0 | 0 | 0.08 | 0.05 | 1 | | | | | |
| PE-Dazzle 594 | 0 | 0.01 | 0 | 0.06 | 0.47 | 0.18 | 0.01 | 0.01 | 0.04 | 0.25 | 0.46 | 1 | | | | |
| PE-Cy5 | 0 | 0 | 0 | 0.01 | 0.21 | 0.3 | 0.03 | 0.03 | 0.01 | 0.68 | 0.12 | 0.44 | 1 | | | |
| PE-Cy7 | 0 | 0 | 0 | 0 | 0.04 | 0.04 | 0.17 | 0.17 | 0 | 0.26 | 0.03 | 0.06 | 0.15 | 1 | | |
| APC | 0 | 0 | 0 | 0.01 | 0.13 | 0.37 | 0.04 | 0.04 | 0 | 0.32 | 0.03 | 0.15 | 0.48 | 0.05 | 1 | |
| APC-Cy7 | 0 | 0 | 0 | 0 | 0.02 | 0.08 | 0.21 | 0.23 | 0 | 0.17 | 0 | 0.02 | 0.09 | 0.29 | 0.2 | 1 |
| --- | BUV395 | Zombie UV | BV421 | BV510 | BV605 | BV650 | BV786 | BV785 | FITC | PerCP-Cy5.5 | PE | PE-Dazzle 594 | PE-Cy5 | PE-Cy7 | APC | APC-Cy7 |

Complexity™ Index: 26.81

**Supplemental Figure 2: The top differentially expressed genes were identified for each cell type identified on the UMAP.**

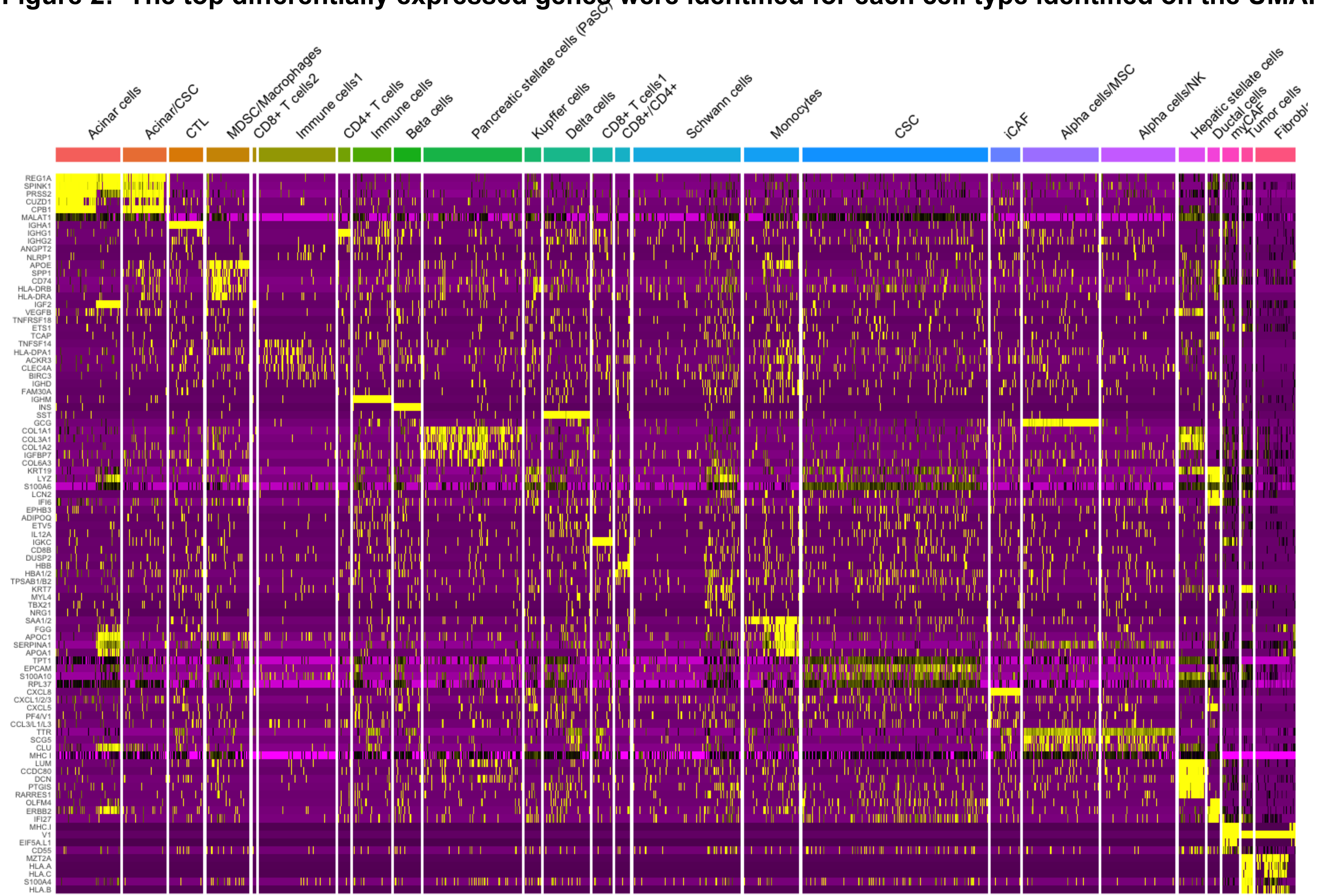

Supplementary Figure 3: The top differentially expressed genes were reported for each cell type identified in tissues analyzed by CosMx whole transcriptomic (WTx) SMI approach.

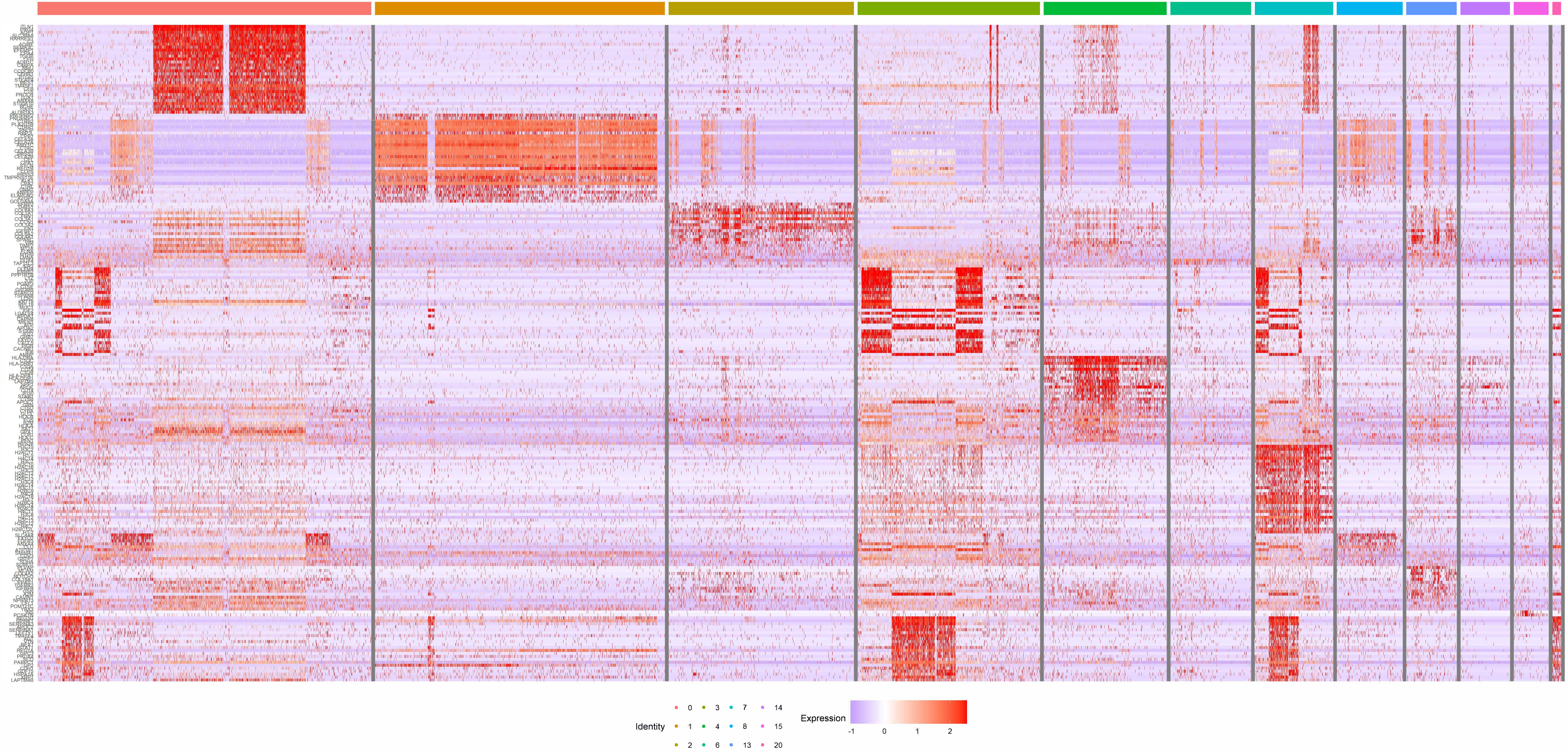

Supplementary Figure 4: The 30 highest expressing genes are reported for adjacent cancer (AC) tissues analyzed by CosMx whole transcriptomic (WTx) SMI.

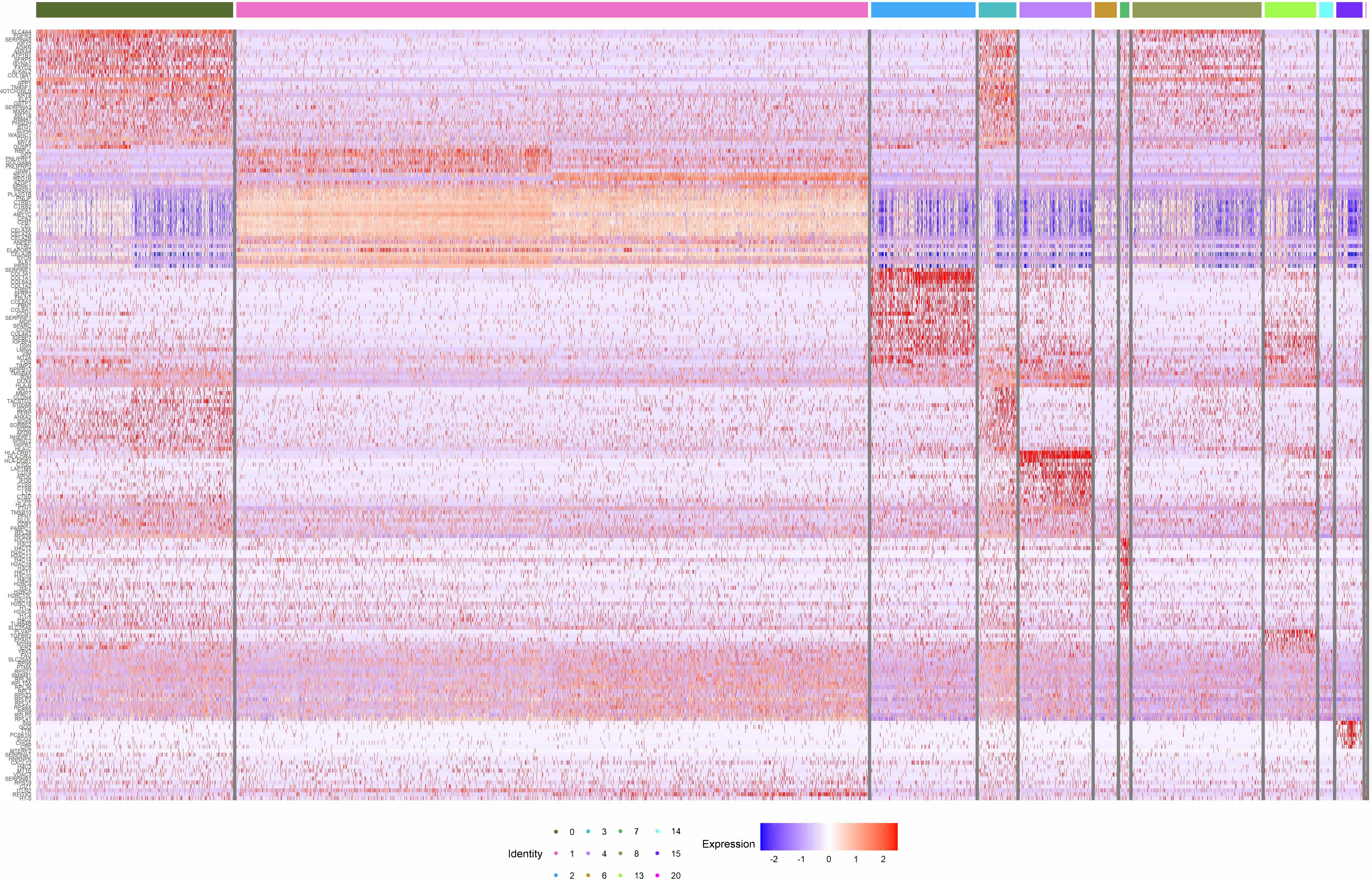

Supplementary Figure 5: The 30 highest expressing genes are reported for IPMN tissues analyzed by CosMx whole transcriptomic (WTx) SMI.

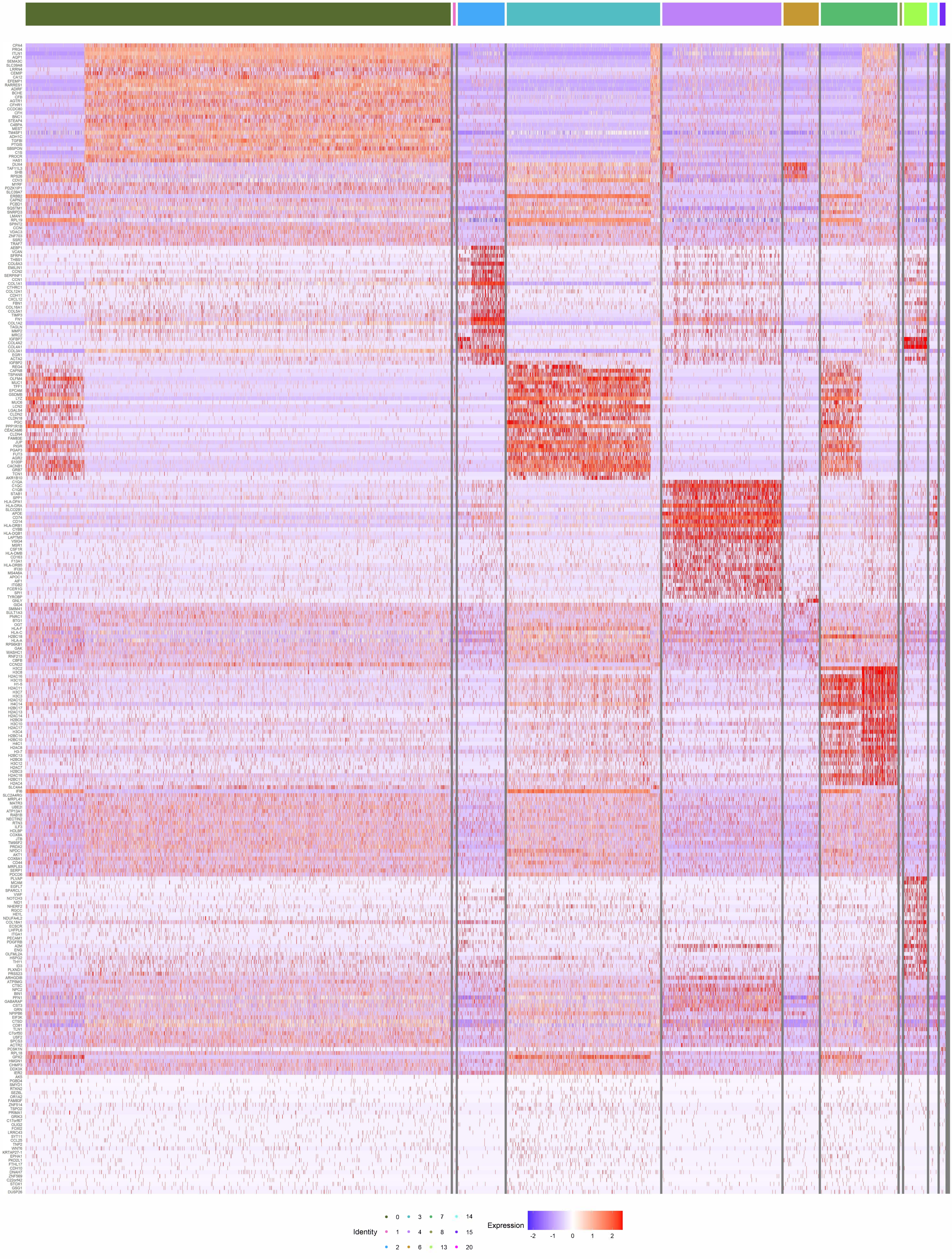

Supplementary Figure 6: The 30 highest expressing genes are reported for PDAC tissues analyzed by CosMx whole transcriptomic (WTx) SMI.

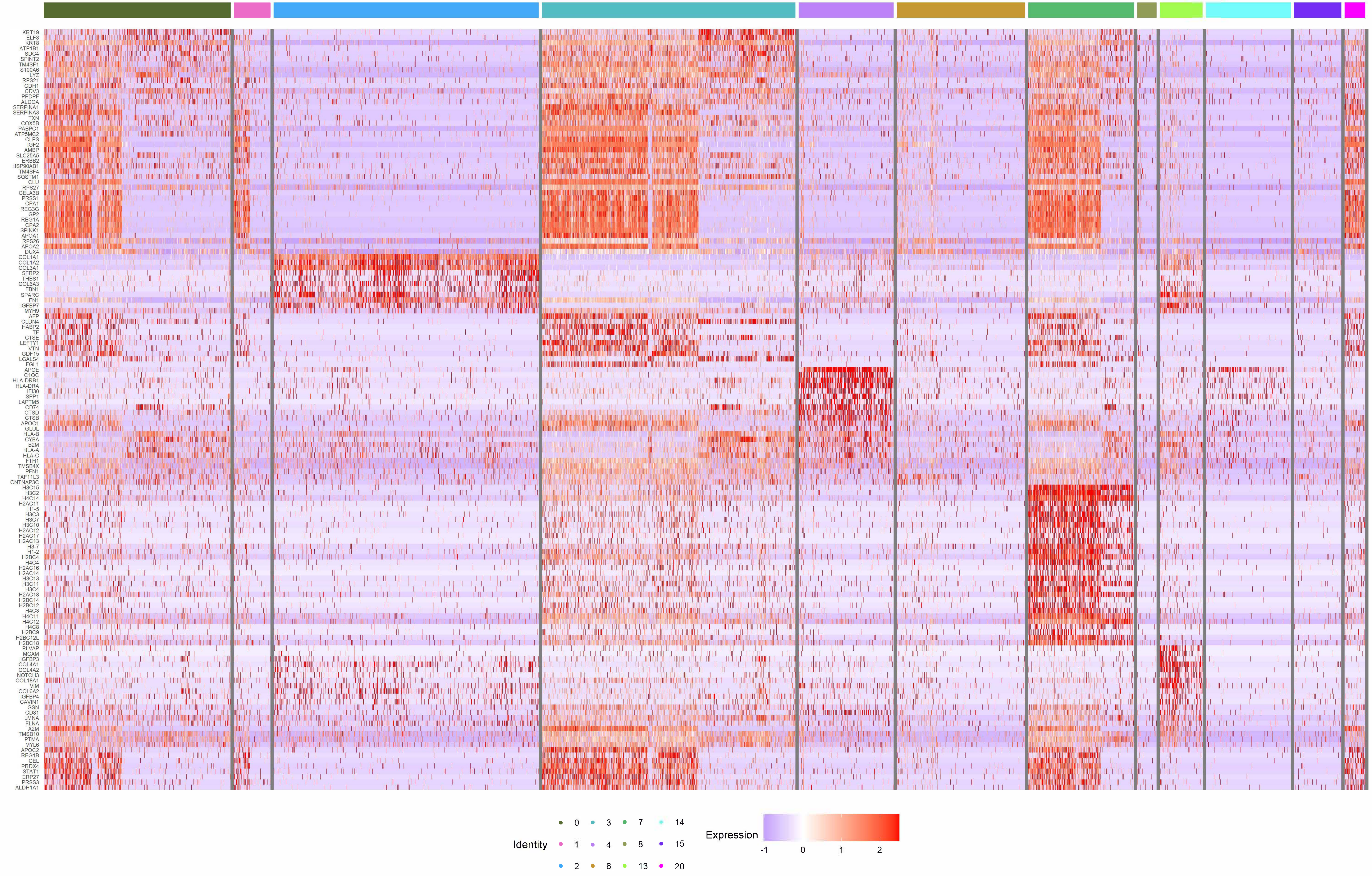

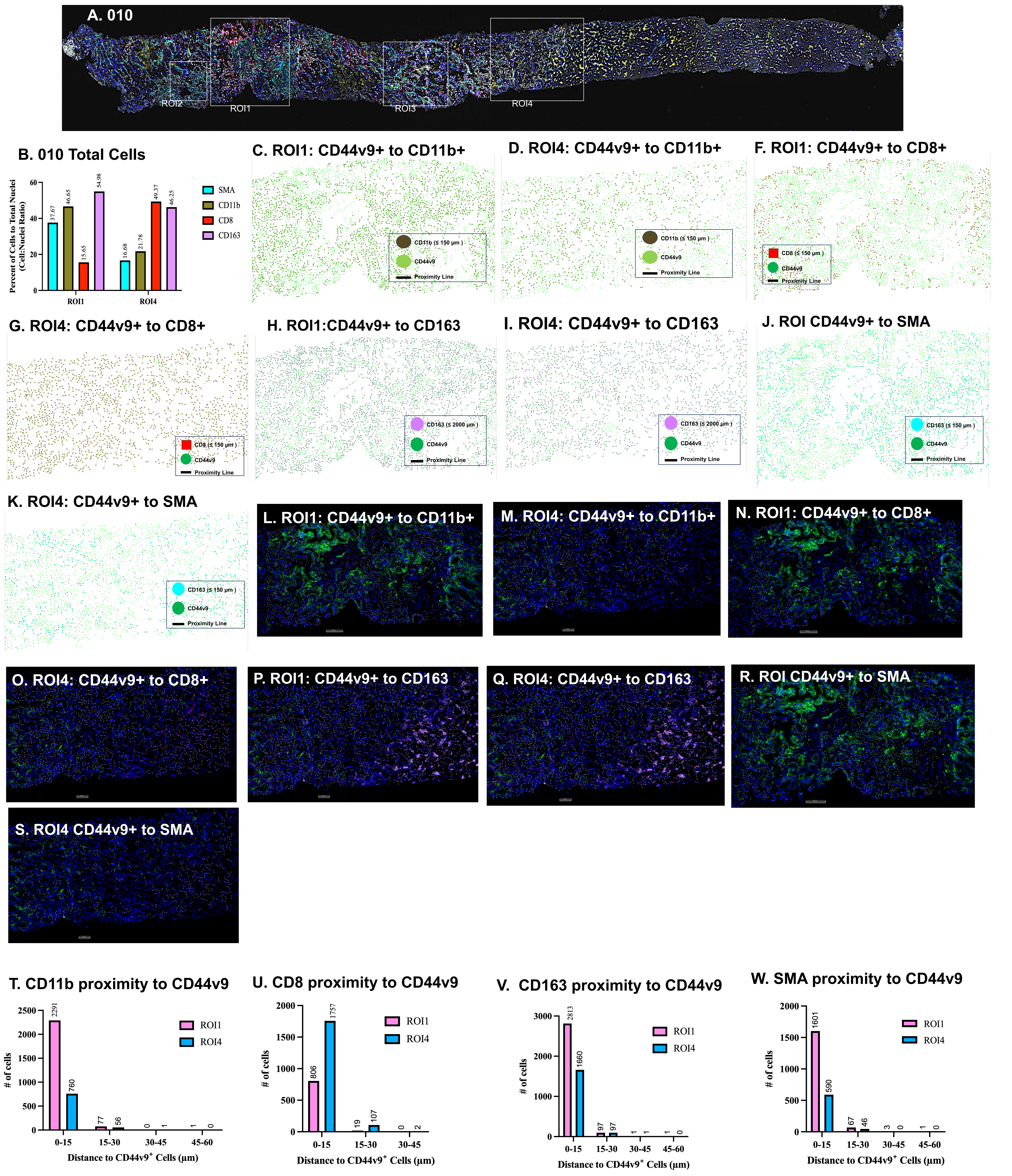
A. 010
ROI2
ROI1
ROI3
ROI4
B. 010 Total Cells
Percent of Cells to Total Nuclei (Cell:Nuclei Ratio)
SMA
CD11b
CD8
CD163
37.67
46.65
15.65
54.98
16.68
21.78
49.37
46.25
ROI1
ROI4
C. ROI1: CD44v9+ to CD11b+
D. ROI4: CD44v9+ to CD11b+
F. ROI1: CD44v9+ to CD8+
CD11b (≤ 150 µm )
CD44v9
Proximity Line
CD8 (≤ 150 µm )
G. ROI4: CD44v9+ to CD8+
H. ROI1:CD44v9+ to CD163
I. ROI4: CD44v9+ to CD163
J. ROI CD44v9+ to SMA
CD163 (≤ 2000 µm )
CD163 (≤ 150 µm )
K. ROI4: CD44v9+ to SMA
L. ROI1: CD44v9+ to CD11b+
M. ROI4: CD44v9+ to CD11b+
N. ROI1: CD44v9+ to CD8+
O. ROI4: CD44v9+ to CD8+
P. ROI1: CD44v9+ to CD163
Q. ROI4: CD44v9+ to CD163
R. ROI CD44v9+ to SMA
S. ROI4 CD44v9+ to SMA
T. CD11b proximity to CD44v9
### of cells
2291
760
77
56
0
1
1
0
ROI1
ROI4
0-15
15-30
30-45
45-60
Distance to CD44v9+ Cells (µm)
U. CD8 proximity to CD44v9
806
1757
19
107
0
2
V. CD163 proximity to CD44v9
2813
1660
97
97
1
1
1
0
W. SMA proximity to CD44v9
1601
590
67
46
3
0
1
0
